# Mapping the allosteric effects that define functional activity of SARS-CoV-2 specific antibodies

**DOI:** 10.1101/2021.12.27.474251

**Authors:** Nikhil K. Tulsian, Palur V. Raghuvamsi, Xinlei Qian, Gu Yue, Bhuvaneshwari D/O Shunmuganathan, Firdaus Samsudin, Wong Yee Hwa, Lin Jianqing, Kiren Purushotorman, Mary M. Kozma, Bei Wang, Julien Lescar, Cheng-I Wang, Ganesh S. Anand, Peter J. Bond, Paul A. MacAry

## Abstract

Previous studies on the structural relationship between human antibodies and SARS-CoV-2 have focused on generating static snapshots of antibody complexes with the Spike trimer. However, antibody-antigen interactions are dynamic, with significant binding-induced allosteric effects on conformations of antibody and its target antigen. In this study, we employ hydrogen-deuterium exchange mass spectrometry, *in vitro* assays, and molecular dynamics simulations to investigate the allosteric perturbations linked to binding events between a group of human antibodies with differential functional activities, and the Spike trimer from SARS-CoV-2. Our investigations have revealed key dynamic features that define weakly or moderately neutralizing antibodies versus those with strong neutralizing activity. These results provide mechanistic insights into the functional modes of human antibodies against COVID-19, and provide a rationale for effective antiviral strategies.

**Teaser:** Different neutralizing antibodies induce site-specific allosteric effects across SARS-CoV-2 Spike protein

## Introduction

A detailed understanding of the biologic determinants that underlie antibody-mediated neutralization of SARS-CoV-2 is critical for informing our development and evaluation of prophylaxes (vaccines) and antibody-mediated therapies for COVID-19 (*1*, *2*). The principal target for development of effective antibody-based antiviral approaches is the viral Spike glycoprotein. Spike proteins are embedded into the viral envelope that bind to host receptors and mediate entry into the host cells. Spike proteins form trimers, with each monomer comprising an N-terminal S1 and a C-terminal S2 subunit separated by S1/S2 cleavage site (*3*–*5*). The S1 subunit of the Spike protein is comprised of an N-terminal domain (NTD), a receptor-binding domain (RBD) and a C-terminal region connecting it to the S2 subunit. The S2 subunit has a multidomain architecture divided into fusion peptide (FP), heptad repeats (HR1 and HR2), central helix (CH), transmembrane region and a C-terminal domain. The three RBDs exist in an equilibrium of “up” or “down” positions (*4*, *6*). In the up-position, the residues that contact with human angiotensin converting enzyme-2 (ACE2) receptor at the surface of the RBD become accessible for binding (*7*, *8*), followed by proteolysis of Spike protein by furin-like proteases at the S1/S2 cleavage site. This promotes TMPRSS2 cleavage and shedding of S1 to release the S2 subunits responsible for membrane fusion (*9*–*11*) with the target cell membrane for entry (*12*, *13*). As the Spike protein serves as the first point-of-contact between the virus and the host, it represents a primary target for neutralizing antibodies against SARS-CoV-2 infection (*14*).

Given the urgent need to curb the pandemic, multifaceted strategies such as mRNA based vaccines (*15*, *16*), small-molecule inhibitors (*17*–*19*), adenovirus-based vaccines (*20*), and antibody-based therapeutics are being employed (*21*–*23*). Neutralizing antibody mediated therapies remain an effective antiviral strategy, as these can be rapidly targeted and/or tested against emerging variants and may also be useful for future Coronavirus-based pandemics (*24*). There are many examples of neutralizing antibodies targeting either Spike NTD or RBD that have been developed for COVID-19 (*25*–*27*), and these mainly interfere with the interactions between SARS-CoV-2 Spike protein with target host receptors such as L-SIGN/DC-SIGN (*28*) and ACE2 (*29*). Antibodies targeting RBD have two major epitopes – (i) ACE2-receptor binding site or the receptor-binding motif (‘RBM’) that is readily accessible (*8*), and (ii) a cryptic epitope that is available only when the RBD is in an ‘up’-position (*30*). In this study, we have characterized a group of novel human antibodies principally derived from convalescent blood samples and describe their dynamic interactions with Spike protein. Using hydrogen-deuterium exchange mass spectrometry (HDXMS), *in vitro* assays, and molecular dynamics (MD) simulations, we have mapped interaction interfaces of antibody-Spike complexes. Earlier studies have primarily focused on Fab or RBD of Spike for antibody characterization, while we have characterized full IgG with the trimeric Spike protein (*1*, *23*, *31*, *32*). These studies offer important insights into the design of antibody cocktails for the most effective neutralization of SARS-CoV-2, with implications against emerging Spike variants of concern.

## Results

### Varying binding affinities and neutralization efficacies of human anti-Spike antibodies

Firstly, we performed biophysical characterization of nine monoclonal human IgG antibodies (‘HuMAbs’), namely LSI-CoVA-014, LSI-CoVA-015, LSI-CoVA-016, LSI-CoVA-017, 4A8 (*31*), 5A6 (*33*), CR3022 (*34*), CoVA-02, and CoVA-39 (*35*). These antibodies were principally discovered from convalescent patients of COVID-19 with the exception of CR3022 that was derived from a SARS-CoV-1 patient and 5A6 which was derived from a human phage-FAB library. The binding activity of each antibody to SARS-CoV-2 Spike trimer and isolated RBD (‘RBD_iso_’) was determined using Quartz Crystal Microbalance (QCM) and enzyme-linked immunosorbent assay (ELISA). As observed by the half-maximal effective concentration (EC_50_) values, seven of the nine antibodies bound strongly to both Spike trimer and RBD_iso_ **(Fig. 1A)**. Antibody LSI-CoVA-017 binds strongly to Spike but not RBD_iso_, suggestive of an epitope outside RBD. Antibody 4A8 binds weakly to Spike and showed negligible binding to RBD_iso_, consistent with previous studies that show 4A8 binds NTD (*36*). Next, we determined the binding kinetics of these HuMAbs against Spike trimer and observed high affinity binding with slow off-rates **(fig. S1, table S1)**. The affinity constants (K_D_) calculated using a 1:1 fit for all HuMAbs were in sub-nM range, with LSI-CoVA-017 being the lowest (0.088 nM). The association-dissociation kinetics clearly indicate stable binding of the HuMAbs with the Spike trimer.

**Figure 1.**
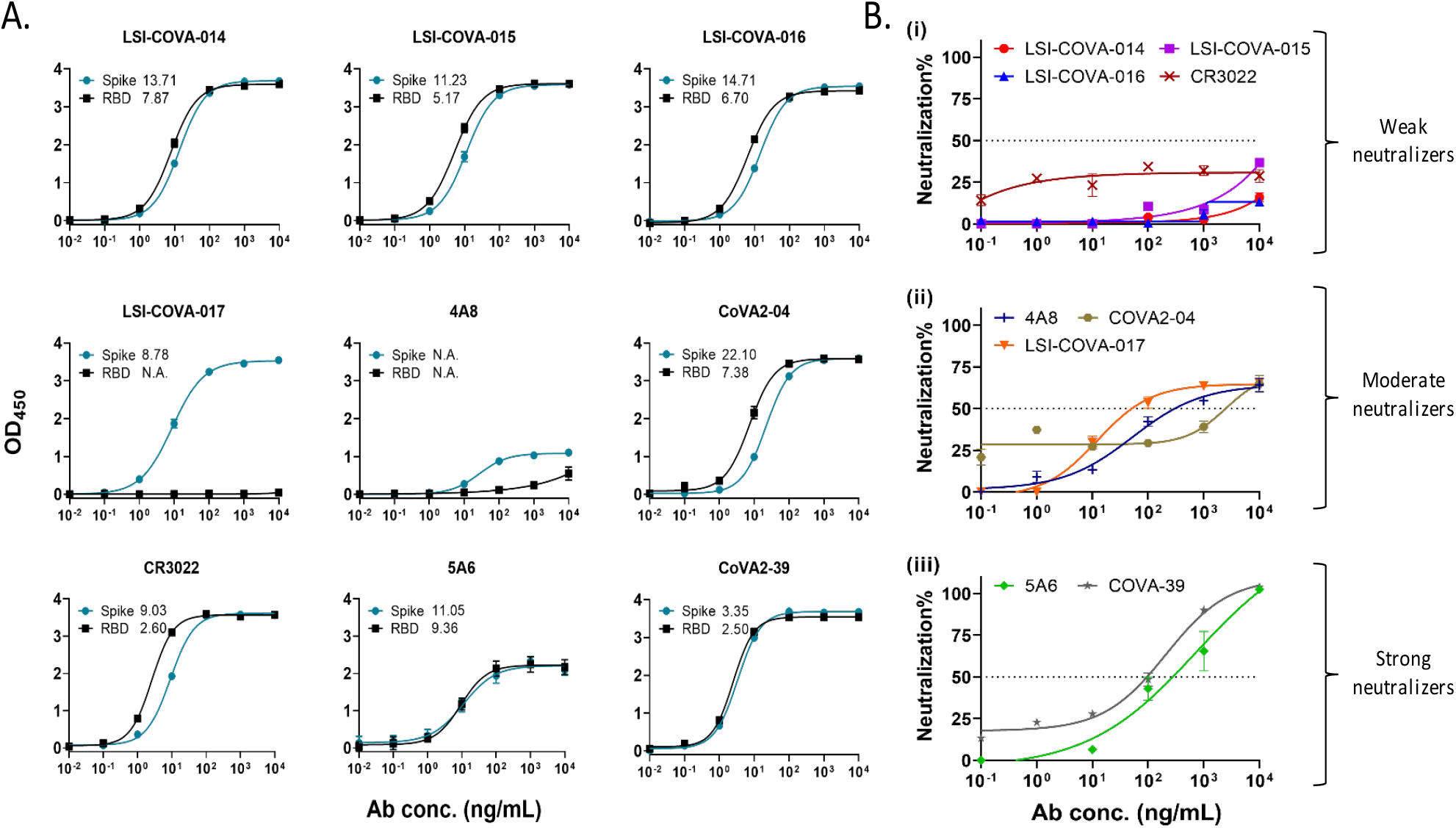
Binding profiles of antibodies against Spike and RBD proteins and their neutralization capacities. (A) Antibody binding activity to Spike and RBD. Nine antibodies at concentrations from 10 pg/mL-10 μg/mL were tested for binding to SARS-CoV-2 Spike (blue plots) and MBP-RBD (black plots) by ELISA, and the EC50 values are indicated. (B) Antibodies at varying concentrations (100 ng/mL-10 μg/mL) were incubated with the pseudovirus lentiviral construct expressing the Spike protein, tagged with luciferase, followed by infection of CHO-ACE2 cells. The chemiluminescence-based luciferase assay readouts were then plotted and presented as percentage neutralization – grouped into (i) weak-, (ii) moderate- and (iii) strong-neutralizing antibodies.

We next investigated their neutralization efficacies using a pseudovirus neutralization test (PVNT). A neutralization capacity of >50% was considered significant. Correspondingly, we observed differential levels of neutralization, wherein LSI-CoVA-014, LSI-CoVA-015, LSI-CoVA-016, and CR3022 showed less than 50% efficacy. On the other hand, 4A8, LSI-CoVA-017, and CoVA2-04 showed significant neutralization capacities, while the highest neutralization was observed for CoVA2-39 and 5A6. On this basis, the antibodies were classified as (i) non-, (ii) weakly-, and (iii) strongly-neutralizing HuMAbs **(Fig. 1B)**.

### Non-neutralizing LSI-CoVA-014, LSI-CoVA-015 and LSI-CoVA-016 bind to cryptic epitope sites on RBD and induce perturbations in Spike trimer structure

The epitopes of the HuMAbs were mapped by comparative HDXMS analysis of complexes with Spike and RBD_iso_. We observed extensive protection in deuterium exchange across peptides spanning RBD of Spike (‘RBD_s_’) and RBD_iso_, in the presence of LSI-CoVA-014, LSI-CoVA-015, LSI-CoVA-016, CR3022, 5A6, CoVA2-04 and CoVA2-39 antibodies. This indicates binding to either RBM or at a site distal to RBM **(Fig. 2, fig. S2)**. Overlapping peptides covering residues 361-395 showed large-scale protection against deuterium exchange in both Spike **(Fig. 2, fig. S2 A, B)** and RBD_iso_ complexes with LSI-CoVA-014, LSI-CoVA-015, and LSI-CoVA-016 HuMAbs **(fig. S2 C, D)**. This region is distal to the RBM site, indicating that these three antibodies bind RBD, without directly interfering with ACE2 receptor binding to Spike protein. In the Spike trimer, the region spanning residues 361-395 becomes accessible only when the RBD adopts an up-position. These differences in deuterium exchange and the identified epitope sites are similar to those observed for CR3022 antibody **(fig. S2D)**, previously characterized as a “cryptic” site binder (*30*). Interestingly, the HDXMS analysis of these three HuMAbs with Spike and RBD_iso_ showed greater deuterium exchange across peptides spanning the RBM **(Fig. 2, fig. S2)**. This observation of increased conformational dynamics at the ACE2 binding site suggests that LSI-CoVA-014, LSI-CoVA-015, and LSI-CoVA-016 binding at the cryptic site induces allosteric destabilization at RBM, which may lead to altered interactions with host receptors.

**Figure 2.**
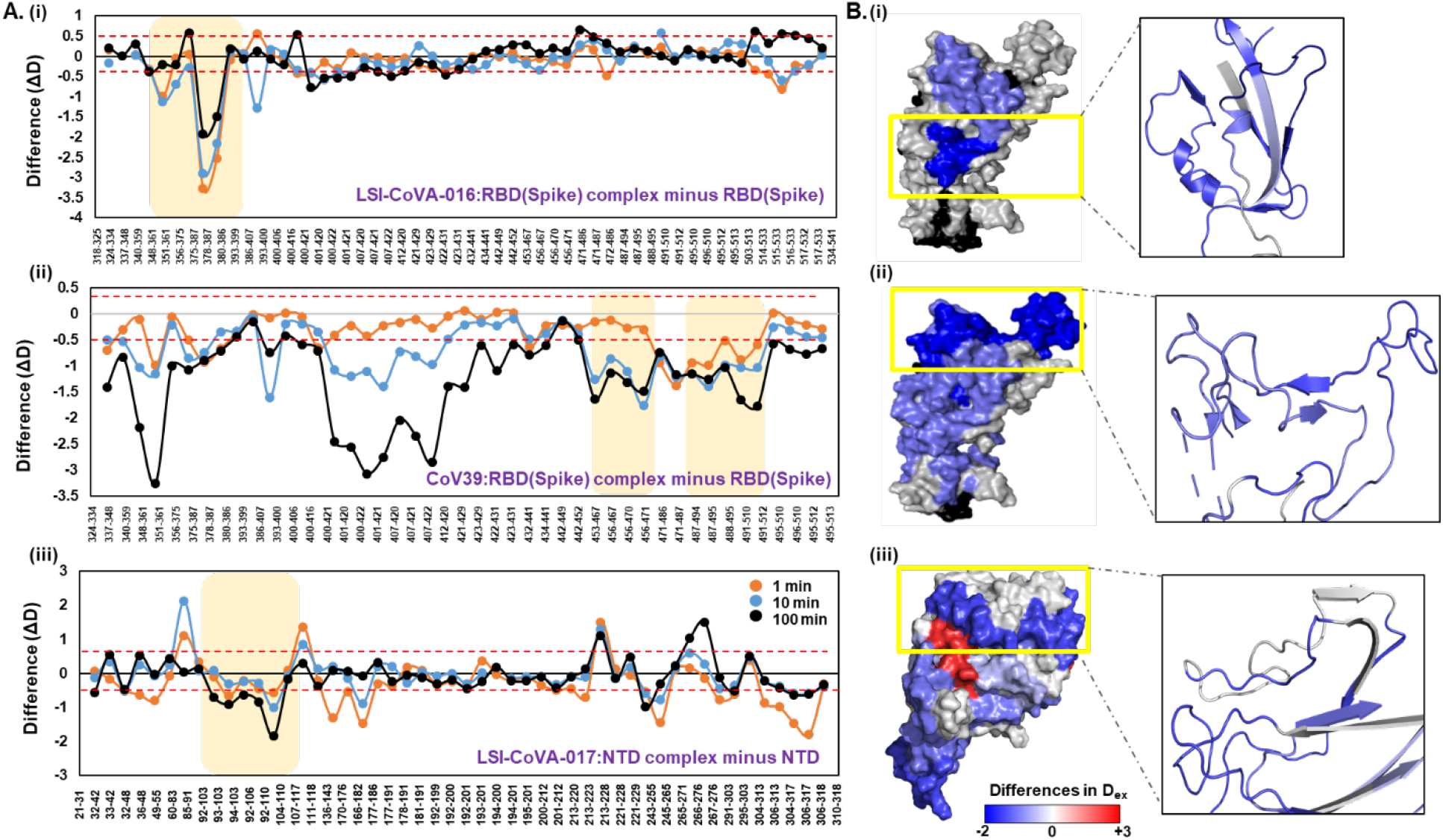
Mapping epitope sites for antibodies with different neutralizing capacities. (A) Plots showing changes in deuterium exchange for (i) non-neutralizing LSI-CoVA-016, (ii) strong-neutralizing CoVA2-39, and (iii) moderate-neutralizing LSI-CoVA-017 antibody complexes with RBD or NTD compared to free antigens, at different labelling times as indicated. Pepsin-proteolyzed fragment peptides are represented by a dot and their residue numbers are indicated (Table S6, Fig. S18). A significant value of ± 0.5 D was considered as threshold and indicated by red-dashed line. Epitope sites are highlighted in yellow. (B) Differences in deuterium exchange values are mapped on to the structure of (i, ii) RBD and (iii) NTD shown in surface representation, as per key. Shades of blue indicate regions with lower deuterium exchange upon antibody binding to the epitope, shown in close-up view.

We then monitored the effect of RBD-binding antibodies on other regions of the Spike trimer. Overall, binding of the non-neutralizing mAbs LSI-CoVA-014, LSI-CoVA-015, and LSI-CoVA-016) to the Spike trimer resulted in similar deuterium exchange kinetics across the S1 **(fig. S3)** and the S2 subunits **(fig. S4)**. While the cryptic epitope sites (361-395) showed similar conformational changes on Spike and RBD_iso_, peptides spanning RBM site exhibited different behavior. Notably, peptides spanning residues 516-533 showed increased deuterium exchange in the presence of antibodies **(Fig. 3A, B, fig. S3)**, confirming that antibody-binding stabilizes RBD in an up-conformation. In this position, many inter-monomer and intra-monomer contacts made by RBD with NTD are broken. In the presence of LSI-CoVA-014, LSI-CoVA-015 or LSI-CoVA-016, an overall increase in deuterium exchange was observed for peptides spanning NTD of Spike trimer **(Fig. 3, fig. S3)**, indicative of greater solvent accessibility or hydrogen-bond disruption as a result of localized destabilization. Increased deuterium exchange was observed at peptides spanning residues 166-182, which interacts with RBD of a neighboring monomer, and residues 289-305 that connects NTD to the central core of the Spike trimer and is indicative of domain movement and disruption of RBD-NTD interactions **(Fig. 3 inset, fig. S3B)**. In addition to these changes at the S1 subunit, most regions of the S2 subunit also showed increased deuterium exchange in the presence of these three mAbs **(Fig. 3A, 3B, fig. S4)**, except the S1/S2 cleavage site that showed decreased deuterium exchange. Prominently, we observed increased deuterium exchange across FP, heptad repeats (HR1 and HR2) and residues 902-916 **(Fig. 3 inset, fig. S4)**, which are all essential for inter-monomer interactions within the Spike trimer. Taken together, the conformational changes observed at the NTD and the S2 subunit suggest antibody-mediated allosteric changes across the Spike trimer, which result in its global destabilization that may lead to partial dissociation of adjacent monomers.

**Figure 3:**
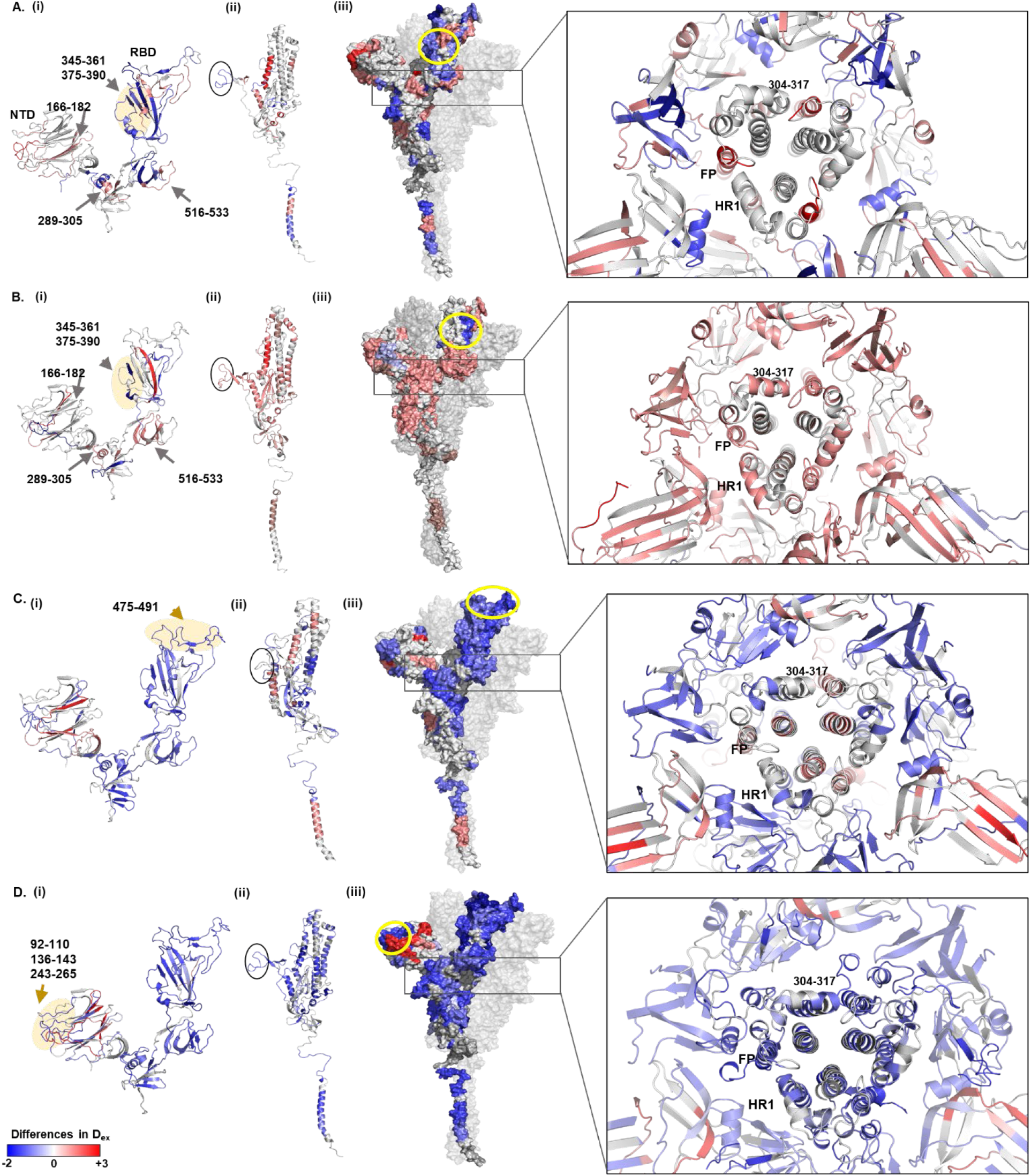
Mechanism of neutralization of Spike trimer by HuMabs. Comparative HDXMS analysis of Spike trimer in the presence and absence of (A) LSI-CoVA-014, (B) LSI-CoVA-016, (C), CoVA2-39, and (D) LSI-CoVA-017 HuMabs were mapped onto the structures of individual (i) S1 and (ii) S2 subunits, and (iii) trimeric Spike protein, as indicated. (i) S1 subunit acts as the primary interface with the neutralizing HuMabs, with the epitope sites highlighted in yellow for (A) LSI-CoVA-014 and (B) LSI-CoVA-016 binding to the cryptic site on RBD, while (C) CoVA2-39 binds at RBM, and (D) LSI-CoVA-017 binds at the NTD. Peptides spanning key regions are highlighted by arrows. Epitopes on RBD (345-361, 375-390, 471-495) and NTD (92-110, 136-143, 243-265) are indicated. (ii) Changes in conformational dynamics of the S2 subunit, upon binding to the antibody, are indicated, with the S1/S2 cleavage site highlighted by black ellipse. (iii) Differences in deuterium exchange are mapped on a monomer of Spike (1-1208 residues), with other two monomers in grey. Epitope sites for the respective antibodies are highlighted by yellow ellipse. Inset highlights a close-up view of Spike trimer along the transverse axis. Differences are mapped onto all the three monomers of Spike. RBD-binding antibodies (LSI-CoVA-014 and LSI-COVA-016) induce destabilizing effects at the inter-protomer contacts, while RBM-binding CoVA2-39 and NTD-binding LSI-CoVA-017 reduce overall conformational dynamics (shades of blue). Peptides spanning residues 304-317, fusion peptide (FP), and heptad repeat 1 (HR1) constituting a part of inter-monomer interaction interface are indicated.

HDXMS analysis of the paratope sites of LSI-CoVA-014, LSI-CoVA-015, and LSI-CoVA-016 antibodies supported these results **(fig. S5)**. Peptides spanning the paratope sites CDRL1-3 and CDRH1-3 showed greater differences in deuterium exchange in the RBD_iso_-bound condition, as compared to Spike-bound antibody complexes **(table S2)**. The epitope sites in RBD_iso_ are not hidden and readily accessible to stably bind the paratope sites. On the other hand, in the Spike trimer, the RBDs move from a down- to an up-position, and antibody binding is further hindered spatially by the NTD and the S2 subunit, leading to less stable antibody binding, compared to RBD_iso_.

### Strong neutralizing HuMAbs bind at RBM and stabilize Spike trimer conformation

Multiple studies have reported high-resolution structures of HuMAbs bound to RBM, including CoVA2-04, 5A6, and CoVA2-39, showing direct competition with ACE2 binding (*37*). However, given that these HuMAbs display various virus-entry neutralization potencies whilst binding to overlapping epitopes, a mechanistic explanation for their contrasting behavior remains elusive, particularly for CovA2-04, a weak neutralizer, as opposed to 5A6 and CoVA2-39 that are strong neutralizers. We therefore monitored the binding of CoVA2-04, 5A6, and CoVA2-39 to the Spike trimer and observed a distinct impact on its conformational dynamics **(Fig. 2, 3)**. A large-magnitude decrease in deuterium exchange was observed for most regions of RBD, particularly the peptide clusters spanning RBM (residues 485-502) of Spike and RBD_iso_ complexes with 5A6, CoVA2-04 and CoVA2-39 **(Fig. 2, fig. S2. fig. S6)**. Interestingly, HDXMS analysis of CoVA2-04 and CoVA2-39 complexes with RBD_iso_ showed lower deuterium exchange predominantly for peptides covering RBM, and only minor changes across other regions of RBD_iso_ **(fig. S2E-G)**. These results indicate that binding of HuMAbs at RBM induces localized changes which lead to a significant reduction in the structural dynamics of RBD, including the peptides spanning the base and linker regions that connect RBD to the Spike trimer. Notably, the Spike variants of concern contain mutations at different sites including E484K-, N501Y- or K417N/E484K/N501 that are localized at RBM, and are reported to reduce the neutralization efficacy of antibodies (*1*, *38*).

HDXMS analysis of Spike complexes with CoVA2-04, 5A6, and CoVA2-39 HuMAbs revealed that they had similar effects on Spike dynamics, wherein an overall reduction in deuterium exchange kinetics was observed. Peptides covering residues 31-42, 92-110, 177-191, 265-276 of NTD showed increased deuterium exchange **(Fig. 3C, fig. S6)**. Some of these peptides span the interface interacting with the RBD or span the C-terminal region of NTD. As binding of these HuMAbs leads to RBD domain movement, it induces NTD movement to disrupt their interface. These changes at NTD are similar to those observed for HuMAbs binding to cryptic epitope sites on RBD **(Fig. 3A,B)**. Significant protection against deuterium exchange was observed at other sites of NTD, and most regions of the S2 subunit. Peptide clusters spanning multiple regions of the S2 subunit, the S1/S2 cleavage site (672-695), regions flanking the fusion peptides (residues 770-782, 878-898, HR1 (927-962), central helix (1003-1031), and residues 1103-1117 showed large-scale decreased deuterium exchange **(Fig. 3C, fig. S6)**. These sites are essential for the Spike trimer to transition from its pre-fusion state to post-fusion state. Taken together, the HDXMS results provide detailed insights into the mechanism of action of these potent neutralizing antibodies CoVA2-04, 5A6, and CoVA2-39, wherein they compete with ACE2 binding, and induce stabilization throughout the Spike trimer to restrict it mobility.

### Recognition of NTD by LSI-CoVA-017 induces global stabilization of Spike trimer

We next investigated the effects of moderate neutralizing HuMAb LSI-CoVA-017 on Spike trimer. In an LSI-CoVA-017-bound state, global protection against deuterium exchange was observed across the Spike trimer **(Fig. 3D)**, with only a few peptides of the NTD showing deprotection **(Fig. 3D, fig. S7A)**. Upon closer examination, peptides spanning residues 92-110, 136-143 and 243-265 **(fig. S7A, bottom panel)** showed large-scale decreased deuterium exchange in the LSI-CoVA-017-bound state. These peptides are positioned towards the outer edge of NTD, and are likely the epitope sites binding to LSI-CoVA-017 **(Fig. 3D, yellow)**. This region also corresponds to the NTD antigenic supersite where all other known NTD-targeting antibodies bind (*25*, *39*). Reduction in deuterium exchange at short labeling times was observed at peptides spanning residues 36-48, 166-182, and 303-318, while increased deuterium exchange was observed for peptides 60-83, 107-117, 213-228, and 266-276 **(fig. S7)**. These differences, mapped onto the structure of NTD **(Fig. 3D, fig. S7A ii)**, revealed that peptides encompassing the epitope site are clustered closely to form a conformational epitope and mediate Spike:LSI-CoVA-017 complexation. These peptides also form the epitopes for previously characterized antibody 4A8, an NTD-binding antibody (*40*). Therefore, we performed HDXMS analysis of 4A8 with Spike trimer and observed large-scale reduction in deuterium exchange at these epitope sites **(fig. S8A)**, akin to stable Spike:4A8 complexation. Thus, LSI-CoVA-017 is an NTD-binding antibody, with both LSI-CoVA-017 and 4A8 being moderate neutralizing HuMAbs.

Large-magnitude decreases in deuterium exchange were observed across all peptides spanning the RBD of Spike bound to LSI-CoVA-017 **(Fig. 3D, fig. S7B bottom panel)**, indicating significantly reduced conformational dynamics across RBD. Peptides (residues 320-350, 516-533) spanning the N- and C-terminal regions of RBD, associated with its domain movement, showed protection against deuterium exchange. This suggests lower solvent accessibility as a result of restricted mobility of RBD in an LSI-CoVA-017-bound state. Similar deuterium exchange patterns were observed for Spike:4A8 complex **(fig. S7)**. To further understand these changes on RBD, we performed HDXMS of LSI-CoVA-017 and 4A8 with RBD_iso_. However, no significant change was observed in deuteration kinetics of RBD_iso_:LSI-CoVA-017 or RBD_iso_:4A8 compared to free RBD_iso_. From these results, it is clear that antibodies binding at the NTD induce allosteric change across RBD. Furthermore, the allosteric effects were propagated to the S2 subunit. Contrary to the changes observed for RBD-binding HuMAbs, decreased deuterium exchange was observed for peptides spanning the S2 subunit in the Spike:LSI-CoVA-017 complex, including the S1/S2 cleavage site **(Fig. 3D, fig. S7C)**. LSI-CoVA-017 binding reduced conformational changes of peptides encompassing FP, central helix, and HR as well **(Fig. 3D iii)**. While both LSI-CoVA-017 and 4A8 binding resulted in similar effects on the Spike trimer, the changes induced by 4A8 HuMAb were less prominent. Overall, these HDXMS results reveal that LSI-CoVA-017 binds at NTD and induces global stabilization of the Spike trimer.

HDXMS analysis of the LSI-CoVA-017 antibody interface showed significant changes across both heavy and light chains in the presence of Spike **(fig. S8A)**. Peptides overlapping CDRH2 (residues 48-70), CDRH3 (residues 96-103), and CDRL2 (residues 48-71) showed protection against deuterium exchange, while CDRL3 (residues 101-129) showed increased deuterium exchange in Spike-bound LSI-CoVA-017 **(table S2)**. Interestingly, similar changes on the paratope sites of 4A8 complexed to Spike were observed **(fig. S8B)**. No significant changes were observed for the light chain of 4A8 with or without Spike, consistent with the high-resolution structures of Spike-4A8 complex (*36*).

The commonalities in effects of 4A8 and LSI-CoVA-017 on Spike, suggests that they have similar modes of neutralization, as reflected by their neutralization capacities. However, our biophysical data showed LSI-CoVA-017 binds Spike trimer with an affinity much greater than 4A8 **(Fig. 1)**. To gain further insights into this, we determined the stoichiometry of Spike:LSI-CoVA-017 complex by size-exclusion chromatography **(fig. S9A, table S3)**. Three chromatographic peaks were detected and analyzed by denaturing polyacrylamide electrophoresis **(fig. S9B)**. Densitometry analysis of different amounts of peak B suggested a binding stoichiometry of three LSI-CoVA-017 per Spike trimer. With a 1:3 stoichiometry of Spike:LSI-CoVA-017, two models are predominant **(fig. S9C)** where: (i) Fab arms from three LSI-CoVA-017 bind to three monomers of Spike; or (ii) two Fab arms of the same LSI-CoVA-017 bind monomers of two different Spike trimers. This is similar to the model predicted for the Spike:4A8 complex (*36*) **(fig. S10)**. Using the most probable model, LSI-CoVA-017 IgG was docked onto one or two Spike trimers. Both binding kinetics as well as detailed characterization of the Spike:LSI-CoVA-017 complex underscores that an LSI-CoVA-017 mAb engages two Spike trimers and each Spike trimer binds to Fab from three different LSI-CoVA-017.

### MD simulations suggest glycan-F_ab_ interaction stabilizes Spike-antibody complexes

To gain high resolution structural insights into the novel antibodies and their interactions with RBD and NTD, we performed docking and MD simulations. Here, Fab domains of each IgG from the current study were modelled and docked onto their respective epitope sites of the Spike protein (RBD or NTD) revealed by our HDXMS, followed by atomistic MD simulations. The rationale here is that the most probably binding pose of Fab to its Spike epitope could be extracted from equilibrated Fab:RBD/NTD MD trajectories, which would then enable us to model full-length Spike:IgG complexes and subsequently infer the avidity and stoichiometry of the antibody. Fab arms of each of LSI-CoVA-014, LSI-CoVA-015, LSI-CoVA-016, and LSI-CoVA-017 HuMAb were modelled and docked onto the epitope sites (RBD and NTD) using HDXMS maps as restraints **(Fig. 4, fig. S10, table S4)**. Multiple orientations of the Fab arms around the defined epitope sites were generated **(fig. S10)**. First, the five top-scoring binding poses were selected and a 200 ns simulation was performed for each. Among the simulated models of Fab:RBD/NTD complexes, multiple models were observed to either displace from epitope site (Model 3 from RBD:LSI-CoVA-014) or completely detach from RBD (Model 4 from RBD:LSI-CoVA-014 and Model 3 from RBD:LSI-CoVA-016) (f**ig. S10**). The trajectories of these displaced or detached Fab:RBD/NTD models were not considered for further analysis. Consequently, we analyzed the trajectories for the remaining four simulated models of the complex. Interestingly, during the simulations, the glycan moiety at N343 of RBD as well as N74, N122 and N149 on NTD were observed to interact with the Fab arm **(Fig. 4A)**. To choose the best model for each antibody, we then measured root means square deviation (RMSD) of the backbone atoms of the Fab domain (**fig. S10**). The most stable model, as indicated by the lowest RMSD, was then selected and two further 200 ns replicate simulations were performed to improve the conformational sampling (Model 2: LSI-COV-014, Model 1: LSI-CoVA-015, Model 4: LSI-CoVA-016, Model 3: LSI-CoVA-017). A stable Fab binding orientation from the most populated cluster was identified by a clustering analysis using concatenated 600 ns long MD trajectories of Fab:RBD/NTD complexes (**Fig. 4A and 4D**). A backbone RMSD cutoff of 0.35 nm of Fab conformations identified 34, 79, 52 and 65 clusters sampled for LSI-CoVA-014, LSI-CoVA-015, LSI-CoVA-016 and LSI-CoVA-017, respectively **(Fig. 4B and fig. S11, S12)**. Contact frequencies calculated between the glycan moieties and Fab from the above triplicate of MD trajectories of each Fab:RBD/NTD complex simulations (as described in Methods) revealed that the glycan interacts with the amino acid residues across all Fab arms of LSI-CoVA-014, LSI-CoVA-015, LSI-CoVA-016 and LSI-CoVA-017 **(Fig. 4B and fig. S13)**. For the amino acid residues interacting with the glycan at N331 and N343, contact frequency maps showed the highest number of contacts and magnitude of contact frequencies made by LSI-CoVA-014 Fab, as compared to those by LSI-CoVA-015 and LSI-CoVA-016 **(Fig. 4B, fig. S13)**. Similarly, our MD trajectories of NTD:LSI-CoV-017 have shown prominent interactions between N74, N122 and N149 glycans and Fab molecule. The contact frequency measured among NTD:LSI-CoV-017 Fab show stable interactions with an even larger interaction surface compared to any of the RBD:Fab complexes (**Fig. 4B and fig. S13**), suggesting a potential role of glycans in forming the antibody epitope.

**Figure 4:**
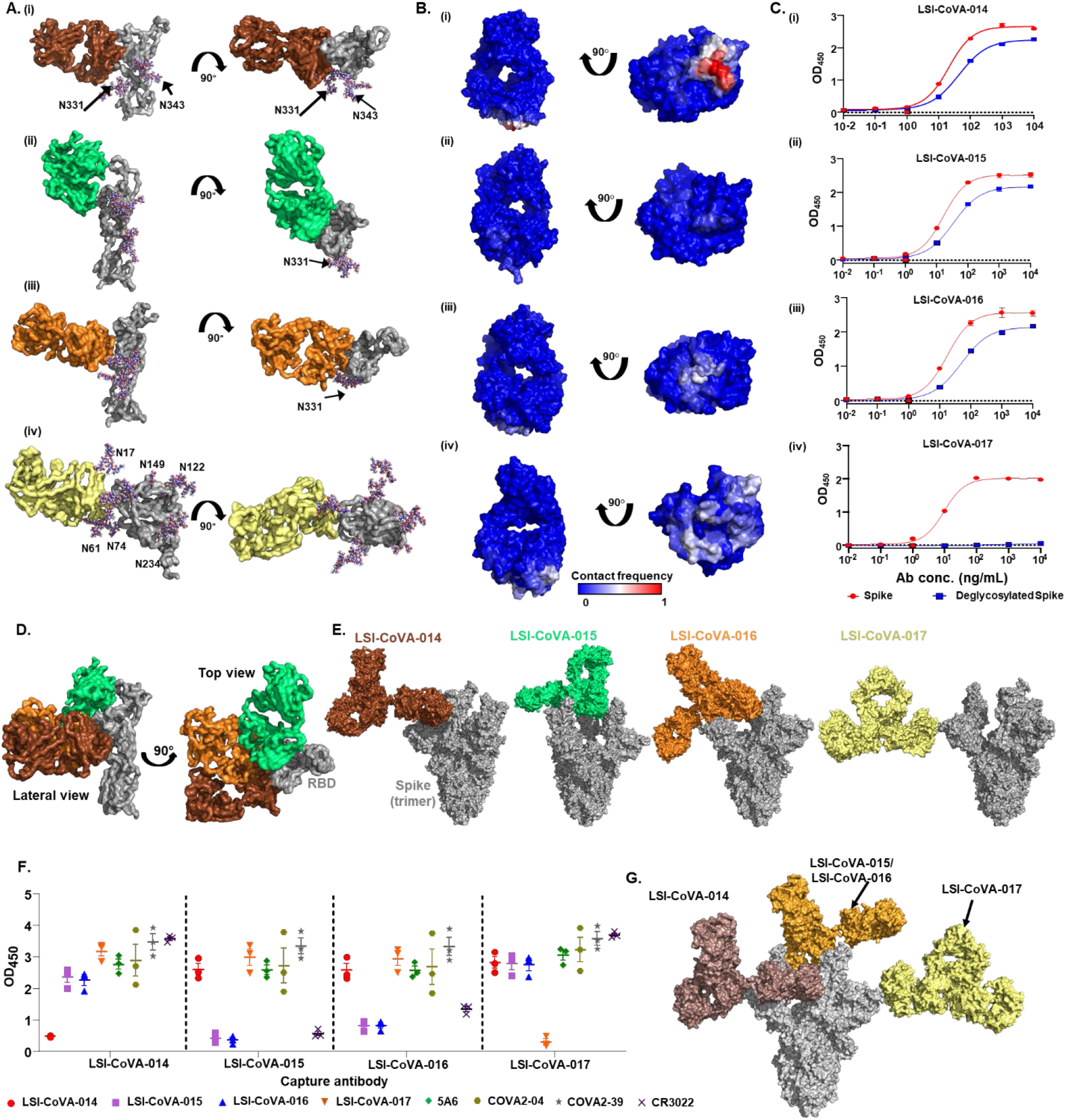
Essential role of glycan groups in facilitating antigen-antibody interactions and determining competitive binding of antibodies to Spike. (A) Representative structures from the most populated cluster (cluster 1) from MD simulation trajectories are depicted in two orientations for: RBD (grey) complexes with Fab of (i) LSI-CoVA-014 (brown), (ii) LSI-CoVA-015 (green), and (iii) LSI-CoVA-016 (orange); and NTD-complex with (iv) LSI-CoVA-017 (yellow). Glycans are shown in ball-and-stick representation. (B) Contact frequencies during simulations between glycan moieties and Fab are mapped onto the structure of each Fab. (C) Plots showing the binding of varying concentrations of (i) LSI-CoVA-014, (ii) LSI-CoVA-015, (iii) LSI-CoVA-016, and (iv) LSI-CoVA-017 antibodies with Spike (red circles) and deglycosylated Spike (blue squares) as determined by ELISA. Average of three individual experiments is represented as mean ± S.E.M. (D) Surface representation (lateral and top views) of Fab LSI-CoVA-014 (brown), LSI-CoVA-015 (green) and LSI-CoVA-016 (orange) with RBD (grey), highlighting their relative orientations. (E) The corresponding full-length IgGs were modelled onto the Spike trimer (grey, PDB: 7A98) based on the position of their Fab as determined by cluster analysis. (F) SARS-CoV-2 Spike (0.1 μg) was captured by LSI-CoVA-014, LSI-CoVA-015, LSI-CoVA-016, or LSI-CoVA-017 using ELISA, detected using peroxidase-labelled monoclonal antibodies. A low OD_450_ value is indicative of impaired binding of the peroxidase-labelled detection antibody, as listed. Data was collected from three individual experiments and represented as mean ± S.E.M. (G) Representation of antibodies binding to different Spike protomers, representative of the competition of LSI-CoVA-015/LSI-CoVA-016 with each other whilst allowing binding of LSI-CoVA-014 or LSI-CoVA-017.

Based upon our HDXMS-guided docked and simulated models, we next studied the binding of four novel HuMAbs with a deglycosylated Spike trimer. Binding of LSI-CoVA-017 was completely abolished upon Spike deglycosylation **(Fig. 4C)**, in contrast to the minor changes observed in binding of LSI-CoVA-014, LSI-CoVA-015 and LSI-CoVA-016 **(Fig. 4C)**, consistent with our simulations showing the most prominent glycan contacts with LSI-CoVA-017. Furthermore, significant reduction of the association-dissociation of these four HuMAbs with varying concentrations of deglycosylated Spike compared to the glycosylated Spike trimer was observed **(fig. S14)**. To validate the significance of deglycosylation to RBD-binding HuMAbs, we tested 5A6 as control, which showed partial reduction **(fig. S14)**. Collectively, these results clearly demonstrate that the LSI-CoVA-017 epitope includes glycan moieties on the Spike protein surface. For other antibodies (LSI-CoVA-014/LSI-CoVA-015/LSI-CoVA-016/5A6) the primary site of binding were non-glycosylated epitopes, as identified by HDXMS with secondary interactions contributed by glycans **(Fig. 4)**. These results provide a view contrary to the prevailing notion that glycans only act as a shield for Spike protein to hide epitope sites from host immune recognition. Our results suggest that non-specific interactions of glycans with the antibodies play a substantial role in stabilizing Fab arm binding at the epitope site.

### Spike:Antibody models and competitive binding provide rationale for combinatorial therapy

A capture ELISA was conducted to evaluate if any of these antibodies compete with each other in binding at the cryptic epitope **(Fig. 4F)** or sterically hinder binding of another antibody to other monomers in the Spike trimer. This allowed us to distinguish the mechanisms of binding and neutralization of RBD-specific antibodies that share the same or highly overlapping epitopes, yet have different affinities and neutralization activities. Competitive binding ELISA results showed similar OD_450_ values between LSI-CoVA-015 and LSI-CoVA-016 as detection or capture antibodies **(Fig. 4F, fig. S15)**. This clearly suggests a significant overlap in binding orientation to the epitopes by LSI-CoVA-015 and LSI-CoVA-016, and is in-line with our HDXMS-constrained docking and MD simulations **(Fig. 4E, fig. S11, S16)**. On the other hand, LSI-CoVA-014 did not prevent binding of LSI-CoVA-015 or LSI-CoVA-016. Docking and MD simulations showed that LSI-CoVA-014 bound to Spike in a different orientation than LSI-CoVA-015 or LSI-CoVA-016, and thus can pair with either LSI-CoVA-015 or LSI-CoVA-016 **(Fig. 4, fig. S16, S17)**. Competition between RBD- and NTD-recognizing antibodies showed that the binding sites for these two antibody classes do not overlap with each other, as observed for LSI-CoVA-014 and LSI-CoVA-017 with LSI-CoVA-015/LSI-CoVA-016 antibodies (**Fig. 4F**).

To infer stoichiometry and plausible mechanisms of neutralization, we then modelled the binding of full-length IgGs to Spike using a representative structure of Fab:RBD/NTD from the previously described clustering analysis performed on concatenated MD simulation trajectories **(fig. S16)**. Modelled full length antibodies showed that RBD-binding antibodies specifically bind to RBD in the open state. Full length LSI-CoVA-015 and LSI-CoVA-016 bound to Spike trimer showed that IgG binding to a single RBD of Spike protomer sterically hinders the binding of a second IgG to the same Spike trimer. In the case of LSI-CoVA-014 and LSI-CoVA-017, the predicted orientation allows the respective full-length antibody to bind all three open RBDs of protomers of the same Spike protein trimer **(Fig. 4G)**. These results are consistent with our competitive ELISA and neutralization assays **(Fig. 4F, fig. S17)**. Additionally, the second Fab arm of LSI-CoVA-017 and LSI-CoVA-014 can bind to a second Spike protein trimer, crosslinking two Spike trimers **(fig. S16)**. Taken together, the Spike:IgG models suggest that the novel antibodies characterized in this study indirectly interfere with ACE2 binding by either crosslinking Spike trimers on the viral surface (LSI-CoVA-014 and LSI-CoVA-017), or by blocking RBD-ACE2 interaction on a single Spike trimer (LSI-CoVA-015 and LSI-CoVA-016).

Overall, our study has identified four novel antibodies isolated from convalescent patients with suboptimal levels of neutralisation efficacy compared to RBM binding antibodies. Considering the mutually exclusive epitope sites complemented by high affinity binding to Spike protein, it would be of interest to explore the use of antibody cocktails against Spike to induce destabilisation into individual monomers or stabilization to reduce the hinge dynamics in order to effectively neutralise the SARS-CoV-2 or its emerging variants. Synergistic effects of selected HuMAbs used in this study were thus evaluated using the PVNT assay. The selected HuMAbs were used in a pairwise cocktail to study the potential synergistic enhancement in neutralization efficacy. When testing a total concentration of 10 μg/mL and in a 1:9 ratio, the paired MAb cocktail of LSI-CoVA-017 and CoVA2-04 displayed a significantly higher percentage neutralization in comparison to the treatment of either single HuMAbs **(fig. S17)**. We did not observe any further addition to the neutralization efficacies of the two potent HuMAbs (CoVA2-39, 5A6) with NTD-binding LSI-CoVA-017. Surprisingly, a combination of NTD- (LSI-CoVA-017) with RBD- (LSI-CoVA-014) antibodies resulted in lower neutralization, than when added alone.

## Discussion

A global effort is currently underway to develop and characterize high-efficacy neutralizing antibodies for SARS-CoV-2 Spike protein and its variants. In this study, we have described antibodies principally derived from convalescent blood of COVID-19 patients and show these can be classified into three groups – non-neutralizers (LSI-CoVA-014, LSI-CoVA-015, LSI-CoVA-016, CR3022), moderately-neutralizing (LSI-CoVA-017, 4A8, CoVA2-04), and strong-neutralizing (CoVA2-39 and 5A6). Importantly, we have described the effects of binding of each of these antibodies to Spike protein and provide mechanistic insights into distinguishing between neutralizing and non-neutralizing antibodies. Antibodies binding to the cryptic site on RBD (LSI-CoVA-014, LSI-CoVA-015, LSI-CoVA-016, and CR3022) alter the localized dynamics at the host ACE2 receptor binding motif and induce increased dynamics at distal sites across the Spike protein. However, this does not result in antibodies with augmented or stronger neutralizing activity. The highest neutralization capacity was observed for 5A6 and CoVA2-39 antibodies, as they bind and modulate Spike dynamics. While RBD has been the primary target for anti-SARS-CoV-2 antibodies, we characterized an NTD-binding HuMAb LSI-CoVA-017 and compared it to a well-characterized HuMAb 4A8. Although these do not block virus attachment to ACE2, they still showed moderate neutralization. The principal mechanism is likely the binding of two Fab arms of LSI-CoVA-017 and 4A8 to neighboring Spike proteins leading to antibody-mediated oligomerization or ‘aggregation’ **(Fig. 4, fig. S9)**. Further, stabilization of Spike trimer and complexation was strengthened by interactions between the glycan groups on Spike with paratope sites of the mAbs.

Determining the simultaneous binding to NTD- and RBD-antibodies denoted that the classes of antibodies are mutually exclusive and allow binding of more than one HuMAb to Spike trimer/SARS-CoV-2 virus. These mechanistic insights into the functional modalities of Spike specific antibodies has important implications for how we evaluate or adapt antibody-based antiviral strategies. The understanding of the conformational dynamics of Spike-antibody complexes allows us to predict the changes and effects that defined HuMAbs will have against new variants of SARS-CoV-2. Consequently, we believe that the findings of this study support the rational design of specific HuMAb cocktails of different classes of antibody to mitigate the risks associated with SARS-CoV-2 mutation as well as to prepare us for new coronavirus pathogens in the future.

## Materials and Methods

### Isolation and cloning of SARS-CoV-2 Spike, RBD specific human antibodies

Memory B cells were isolated from Peripheral blood mononuclear cells (PBMCs) derived from blood samples from COVID-19 convalescent patients using a Human Memory B cell isolation kit (Miltenyi Biotec, GE). Small pools of purified Memory B cells were seeded into 384-well plates on irradiated CD40L-expressing feeder cells for differentiation into plasma cells as described previously (*41*). After 7 days of culture, supernatants from B cell pools were screened for binding activity on SARS-CoV-2 Spike by ELISA. Antibody Heavy and Light Chain variable regions were cloned from positive wells by PCR using Collibri^™^ Stranded RNA Library Prep Kit for Illumina^™^ Systems (Thermo Fisher, SG) and whole human IgG reconstructed as described previously (*42*). Confirmation of binding specificity of cloned human monoclonal antibodies was confirmed by ELISA.

### Antibody and antigen expression and purification

Genes coding for variable regions of antibody heavy and light chain were synthesized and cloned into vector pTT5 expression vector (National Research Council Canada, NRCC) by Twist bioscience. Antibody heavy and light chain constructs were transfected into HEK293-6E mammalian expression cells at a concentration of 400 ng/ml in Branched-PEI (Polyethylenimine)/150 mM NaCl. 7 days post-transfection, culture supernatant containing the expressed antibodies was harvested via centrifugation, filtered through a 0.22 μm filter unit (Merck, SG) and loaded onto a MabSelect Sure column (Cytiva, SG) to purify the antibody. The purified antibody was subjected to buffer exchange using Vivaspin centrifugal concentrator (Sartious, GE) and concentrated in 1x PBS, pH 7.2.

A near full-length Spike (S) protein, excluding transmembrane domain and cytoplasmic tail, of SARS-CoV-2 (16-1208; Wuhan-Hu-1; GenBank: QHD43416.1) was codon optimized for insect cell expression and cloned into pfastbac expression vector (Biobasic, SG). Gene was followed by a HRV 3C site, 8x-Histidine tag, and a streptavidin tag at the C-terminus. A double mutant Spike construct was generated by mutating RRAR (682-685) into DDDDK and residues KV (986-987) into PP. SARS-CoV-2-Spike construct was expressed in Spodoptera frugiperda Sf9 cells following instructions from bac-to-bac baculovirus expression system (Thermo Fisher, SG). Briefly, bacmid of SARS-CoV-2 Spike was generated, purified and used for transfection using cellfectin II (Thermo Fisher, SG). Viral stocks obtained from transfection were amplified and used for protein expression. Culture supernatant was harvested by centrifugation at day 4 post infection. Spike protein was affinity purified using a 5mL HisTrap excel column (Cytiva, SG). Purified Spike was concentrated using a 100 kDa cut-off concentrator (Sartorius, GE) and loaded onto a HiLoad 16/60 superdex 200 pre-equilibrated with 20 mM Hepes, pH 7.5, 300 mM NaCl, 5% glycerol. Peak fractions corresponding to the trimer size of Spike protein were collected and concentrated. SARS-CoV-2-RBD was expressed and purified as described previously(9). Plasmid Sars-Cov-2 Spike HexaPro (addgene) was used to transfect HEK 293 cells with polyethylenimine (PEI). Culture supernatant was harvested on day 7 post transfection. Spike hexapro was purified following the same protocol as described for spike construct expressed by Sf9 cells.

The purity and integrity of the Spike, RBD and the antibodies were determined by denaturing polyacrylamide electrophoresis.

### Antibody binding activity analysis using enzyme-linked immunosorbent assay (ELISA)

Spike was coated at 100 ng/well and RBD at 200 ng/well onto 96-well flat-bottom maxisorp binding immunoplates (SPL Life Sciences, SG) and incubated overnight at 4 °C. The following steps were conducted at room temperature. Plates were washed three times in PBST (phosphate buffer with 0.05% Tween 20) and blocked with 350 μL of blocking buffer (4% skimmed milk in PBST) for 90 min. A 10-fold serial dilution was performed for each primary antibody to get seven concentrations ranging from 10 μg/mL to 0.01 ng/mL. Plates wash step was repeated and the diluted primary antibodies were added at 100 μL/well. Reference wells with no primary antibody (blocking buffer only) were included. After 60 min incubation, plate wash step was repeated and the detection antibody, goat anti-human IgG-HRP (Thermo Fisher, SG) was added at 100 μL/well for an incubation of 60 min in dark. This was followed by the plate wash step and an incubation with 1-Step Ultra TMB-ELISA (Thermo Scientific, SG) at 100 μL/well for 3 min. Reaction was stopped with 100 μL 1M H2SO4 per well. Optical density at 450 nm was measured using a microplate reader (Tecan Sunrise, SG). Each antibody was tested in three independent replicates and their average values were used to generate the plots.

### Biophysical characterization of binding of mAbs to Spike

Trimeric Spike or PNGase F treated Spike HexaPro (0.03 μM in 10 mM sodium acetate buffer, pH 4.5) was immobilized onto the surface of a LNB-Carboxyl quartz crystal chip (Attana, SE) using sulfo-NHS/EDC. Unbound carboxyl groups were quenched using ethanolamine. A reference chip was similarly activated and quenched, but with no protein bound to its surface. Chips were stabilized in the Attana Quartz Crystal Microbalance (QCM) instrument and experiments performed at 22 °C using a flow rate of either 20 μL/min or 25 μL/min. Running buffer was 10 mM HEPES, 150 mM NaCl, 0.005% Tween 20, pH 7.4 (HBST). Injections of 35 μL, were made in triplicate over both chips, and both association and dissociation phases of each interaction were recorded for 380 seconds using Attana software. Experiments were performed on a variety of ligand densities to control for avidity. Reference curves were subtracted using Attester software and the resulting curves imported into TraceDrawer (Ridgeview Instruments) for kinetics evaluation.

### SARS-CoV-2 pseudotyped lentivirus production

Pseudotyped viral particles expressing SARS-CoV-2 Spike proteins were produced using a third-generation lentivirus system. A reverse transfection methodology was used for this assay. At Day 1, 36 ×10^6^ HEK293T cells were transfected with 27 μg pMDLg/pRRE (Addgene, US), 13.5 μg pRSV-Rev (Addgene, US), 27μg pTT5LnX-WHCoV-St19 (SARS-CoV2 Spike) and 54 μg pHIV-Luc-ZsGreen (Addgene, US) using Lipofectamine 3000 transfection reagent (Thermo Fisher, SG) and cultured in a 37 °C, 5% CO2 incubator. On Day 4, supernatant containing the pseudoviral particles was harvested and filtered through a 0.45 μm filter unit (Merck, SG). The filtered pseudovirus supernatant was concentrated using 40% PEG 6000 by centrifugation at 1600 g for 60 min at 4 °C. Lenti-X p24 rapid titre kit (Takara Bio, JP) was used to quantify the viral titres, as per manufacturer’s protocol.

### Pseudovirus neutralization assay (PVNT)

On Day 0, CHO cell lines with stable expression of ACE2 were seeded at a density of 5 x104 cells in 100 μL of complete medium [DMEM/high glucose with sodium pyruvate (Thermo Fisher, SG), supplemented with 10% FBS (Thermo Fisher, SG), 10% MEM non-essential amino acids (Thermo Fisher, SG), 10% geneticin (Thermo Fisher, SG) and 10% penicillin/streptomycin (Thermo Fisher, SG)] in 96-well white flat-clear bottom plates (Corning, US). Cells were cultured in 37 °C with humidified atmosphere at 5% CO2 for one day. The next day, the respective monoclonal antibodies (mAbs) were serially diluted 10 times in sterile 1x PBS. The diluted samples were incubated with an equal volume of pseudovirus to achieve a total volume of 50 μL, at 37 °C for 1 h. The pseudovirus-antibody mixture was added to the CHO-ACE2 monolayer cells and left incubated for 1 h to allow pseudotyped viral infection. Subsequently, 150 μL of complete medium was added to each well for a further incubation of 48 h. The cells were washed twice with sterile PBS. 100 μL of ONE-gloTM EX luciferase assay reagent (Promega, SG) was added to each well and the luminescence values were read on the Tecan Spark 100M. The percentage neutralization was calculated as follows:

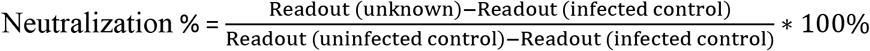

### Neutralization by Antibody pairs

Pair-mAb cocktail in the ratio of 1: 9 to a final total concentration of 10 μg/mL, or single huMAb at a concentration of 10 μg/mL were incubated with pseudovirus lentiviral construct expressing the SARS-CoV-2 Spike protein. The antibody:pseudovirus mixtures were then added to CHO-ACE2 cells. The chemiluminescence readout from the luciferase-tagged reporter in the lentiviral construct, was then plotted and represented as percentage neutralization. The assay was conducted in triplicates. The data is shown as mean ± s.d. GraphPad Prism was used for plots and one-way ANOVA statistical analysis.

### Determining stoichiometry of LSI-CoVA-017 binding to Spike

80 μg spike protein and 100 μg LSI-CoVA-017 (1:1.25 molar ratio) were mixed and incubated at room temperature for 30 min, prior to injection on a Superose 6 Increase 10/300 GL column (GE Healthcare, SG) in buffer (20 mM Tris, 200 mM NaCl, pH 8). The resulting chromatographic eluted peaks were analyzed by SDS-PAGE. Densitometry analysis was carried out using Image Lab (v6.1.0 build 7, Bio-Rad, SG). Briefly, 1-4 μg of Spike and LSI-CoVA-017 were loaded on SDS-PAGE as quantification standards. 40 μL and 25 μL of peak A, and 100 μL and 50 μL of peak B were loaded on SDS-PAGE for quantification. The absolute quantities of Spike, heavy and light chains of LSI-CoVA-017 were estimated based the band intensities relative to the quantification standards. For each lane, the quantities of LSI-CoVA-017 were then derived from the quantities of either heavy chain or light chain. Both methods estimated similar quantities of LSI-CoVA-017, and thus consistently suggest a binding stoichiometry of three LSI-CoVA-017 bound per Spike trimer. In addition, consistent results of the two different loading amounts per peak A or peak B provided further confidence in the reliability of the densitometry. (Refer to Table S1 for detailed calculation).

### Hydrogen-Deuterium Exchange Mass Spectrometry (HDXMS)

#### Deuterium labeling of free proteins

Purified Spike trimer (8 μM) and RBD_iso_ (67 μM), solubilized in aqueous PBS (pH 7.4) were diluted in 20x deuterated PBS to attain 90 % final D2O concentration (*11*). The deuterium labelling was performed for 1, 10, 100 min for Spike protein, and 1 and 10 min of RBD_iso_ respectively. Similarly, HDX was performed for nine convalescent antibodies - LSI-CoVA-014, LSI-CoVA-015, LSI-CoVA-016, and LSI-CoVA-017 identified in this study; and CR3022, CoVA2-04, CoVA-39, 4A8, and 5A6 (previous studies). All deuteration reactions were performed at 37 °C.

#### Deuterium labelling of antigen-antibody complexes

For HDXMS of antibody-protein complexes, saturating concentrations of antibody to the antigen were used. For each antigen-binding site on huMAb was considered as one antigenic site, and mixed Spike trimer in 1:3 (protein:antibody), and 1:1 RBD-antibody stoichiometry. As the antibodies bind with a high affinity, the antigen-antibody complexes were incubated for 30 min at 37 °C to achieve >90% binding, before each deuterium exchange reaction. The resultant protein:antibody complexes (~100 pmol) were diluted in 20x deuterated PBS (90% final concentration of D2O) and subjected to 1, 10, and 100 min hydrogen-deuterium exchange timescales (Table S5).

#### LCMS Analysis

At the end of labeling times, each deuteration reaction was quenched by adding pre-chilled quench solution (1.5 M Gn-HCl, 0.25 M TCEP-HCl) to lower the pHread to ~2.5 and incubated at 4 °C on ice for 1 min. The quenched samples were injected onto nanoUPLC™ HDX sample manager (Waters, USA) for proteolysis by immobilized pepsin cartridge (Enzymate BEH pepsin column 2.1 × 30 mm (Waters, USA)), with a continuous flow of 0.1% formic acid in water (UPLC grade, Merck, GE) at 100 μl/min. The pepsin proteolyzed peptides were trapped on a 2.1 × 5 mm C18 trap (ACQUITY BEH C18 VanGuard Pre-column, 1.7 μm, Waters, USA) and then eluted by a gradient of 8-40% of 0.1% formic acid in acetonitrile using reverse phase column (ACQUITY UPLC BEH C18 Column, 1.0 × 100 mm, 1.7 μm) pumped at a flow rate of 40 μl/min by nanoACQUITY binary solvent manager (Waters, USA). Peptides were ionised using electrospray ionisation method and sprayed onto Synapt G2-Si mass spectrometer (Waters, UK) as described previously (*11*).

#### Data Processing

Protein Lynx Global Server (PLGS) v3.0 was used to identify the detected peptides from mass spectra of undeuterated samples. A separate sequence database of each protein along with the its respective purification tags were used to identify pepsin proteolyzed peptides, as described previously (*11*). Identified peptides with mass tolerance of <10 ppm were analyzed for deuterium uptake using DynamX v3.0 (Waters, USA). All deuterium exchange experiments were performed in triplicate and reported values are not corrected for deuterium back exchange (Table S5-S7).

### Homology modelling of single Fab arm

Homology models of Fab arms of LSI-CoVA-014, LSI-CoVA-015, LSI-CoVA-016, and LSI-CoVA-017 were modelled using Modeller version 9.21 (*43*). Position specific iterative - BLAST was used to identify the template structures with high sequence identity and structures available at PDB were chosen. 7K8R, 6DWZ, 6DF2, and 7JXE were used as template structures to model peptides spanning heavy chain and light chain of LSI-CoVA-014, LSI-CoVA-015, LSI-CoVA-016, and LSI-CoVA-017 respectively. Heavy (1-226) and light (1-216) chains were modelled separately along with intramolecular disulphide bonds. For each antibody, 100 models were generated, and the model with lowest discreet optimized energy (DOPE) score was chosen to generate a Fab arm. Hetero dimers containing heavy and light chain for respective antibodies were complexed by aligning to the respective template structures in PyMol (Schrodiner Inc, USA). Corresponding full-length IgGs were modelled using Swiss Modeller server using 1IGT and 1MCO structures as templates (sequence identity > 90%).

### Glycosylated RBD, NTD modelling and antigen-Fab complex generation

Atomic coordinates of RBD and NTD from full-length Spike protein (PDB: 6XR8 (*44*)) was isolated and further modelled. Glycan chains were constructed using CHARMM-GUI (*45*) glycan reader and Modeller (*46*) at the reported glycosylation sites i.e., N331 and N343 of RBD, and N17, N61, N74, N122, N149, N165, N234, and N282 of NTD (*5*, *47*). The glycan composition reported by previous studies was used to model at the respective glycosylation sites (*5*). The epitope sites on RBD and NTD identified using HDXMS for each antibody were chosen to perform a biased docking using ClusPro 2.0 webserver (*48*). The peptides showing protection in the presence of antibody were provided as input under attractive amino acid residues at the protein-protein interaction interface in ClusPro, using a specific module for antigen-antibody docking. A total of 10 different poses or orientations of Fab bound to RBD/NTD were generated at the epitope site involving complementarity determining regions (CDR) of Fab arms. Each pose was inspected using VMD (*49*) and the top five poses with highest docking score were chosen to perform atomistic MD simulations from each Fab:RBD/NTD complex.

### Simulation setup and protocol

Atomistic MD simulations were performed to identify a structurally stable orientation of Fab bound to RBD and NTD. A total of 20 Fab:RBD/NTD complexes from four different Fab arms were modelled using CHARMM-GUI and simulated in GROMACS package version 2018.4 (*50*). Each system was parameterized using CHARMM36m force field (*51*) and system was solvated in TIP3P water molecules with 0.15 M NaCl added to neutralize the system. The heavy atoms in the Fab:RBD/NTD complex were restrained using a force constant of 1000 kJ mol-1 nm-1 to perform a 100 ps long equilibration of the system. 310 K temperature was maintained using the Nóse-Hoover thermostat with 1.0 ps time constant and 1 atm pressure was kept by isotropic coupling to the Parrinello-Rahman barostat with time constant of 5.0 ps (*52*, *53*). Electrostatic interactions were calculated using the smooth particle mesh Ewald’s method with a real-space cut-off of 1.2 nm. The Van der Waal’s interactions were truncated at a distance of 1.2 nm with a force switch smoothing function imposed from 1.0 to 1.2 nm. LINCS algorithm (*54*) was used to constrain all the covalent bonds with an integration time step of 2 fs. A 200 ns long production run was performed for all the 20 models. The best model for each of the four Fab arms were then selected based on the lowest backbone root means square deviations (RMSDs). Two further 200 ns independent repeat simulations with different starting velocities were performed for these models. The list of simulations performed are provided in Table S4.

Most of the analysis of trajectories were performed using GROMACS analysis package. Pair-wise distances between atoms of glycan and Fab molecule were calculated and any pairwise distance less than 0.4 nm was recorded as an interaction point. The ratio between number of frames the pair-wise distance recorded to be <0.4 nm and total number of frames in each simulation i.e., 2000 is reported as the contact frequency between glycan and Fab molecule. Further, to identify the stable or dominant orientation of Fab bound to RBD and NTD, we performed a cluster analysis using the GROMOS methods and a RMSD cut-off of 0.35 nm. The central structure from the most populated cluster was chosen to perform further analysis and modelling. The trajectories of models 3 and 4 from RBD_iso_: LSI-CoVA-014 and model 4 and model 3 from RBD:LSI-CoVA-016 were not considered for cluster analysis.

### Capture ELISA

Monoclonal human IgG1 antibodies LSI-CoVA-014, LSI-CoVA-015, LSI-CoVA-016, and LSI-CoVA-017 were conjugated individually to peroxidase as per manufacturer’s protocol (Abnova, TW). The final concentrations of conjugated antibodies are 1 mg/mL. Unconjugated LSI-CoVA-014, LSI-CoVA-015, LSI-CoVA-016, and LSI-CoVA-017 were diluted in PBS to a final concentration of 10 μg/mL, and coated on 96-well maxisorp binding immunoplates at 100 μL/well for overnight incubation at 4 °C. Plate wash and blocking steps were conducted as mentioned above. SARS-CoV-2 Spike protein at 1 μg/mL diluted in blocking buffer was added to each well, 100 μL/well for 1 h incubation. Plate wash step was repeated. Peroxidase-conjugated antibodies were added individually at a final concentration of 1 μg/mL (diluted in blocking buffer) for 1 h incubation protected from light. All following steps were performed as mentioned above.

### Binding activity of antibodies to PNGase F treated Spike

20 μg of SARS-CoV-2 Spike trimer was deglycosylated by incubating with 2.5 μL of PNGase F (NEB, SG) under native condition at 37 °C for 4 h. Spike trimer and deglycosylated Spike trimer were coated at 1 μg/ml, 100 μL/well on 96-well maxisorp binding immunoplates for 1 h at room temperature. Following steps of the ELISA are the same as described above. Monoclonal antibodies LSI-CoVA-014, LSI-CoVA-015, LSI-CoVA-016 and LSI-CoVA-017 were tested at several concentrations ranging from 0.01 ng/mL to 10 μg/mL.

### Donor recruitment

Blood samples were collected in accordance with the Helsinki declaration after written consent from volunteers under an approved Institutional Review Board protocol NUS-IRB REF NO: LH-20-013.

## Acknowledgments

We would like to thank Ms. Eve Zi Xian Ngoh for her contribution in discovery and production of 5A6.

## Author contributions

Conceptualization: NKT, PVR, PJB, GSA, PAM

Methodology: NKT, PVR, QX, GY, BS, FS, LJ, WYH, BW, MMK, KP

Investigation: NKT, PVR, MMK, JL, CW, PJB, GSA, PAM Visualization: NKT, PVR, XQ, GY, LJ

Funding acquisition: PAM, PJB

Project administration: MMK, PAM Supervision: NKT, JL, CW, PJB, PAM

Writing – original draft: NKT, PVR, XQ, LJ

Writing – review & editing: NKT, PVR, XQ, FS, JL, PJB, GSA, PAM

## Data and materials availability

All data, code, and materials used in the analysis is available upon request. Request of materials would be done upon signing a material transfer agreement (MTA) with PAM. Request for code for molecular dynamics simulations should be to PJB, while other resources and mass spectrometry related to NKT. All data are available in the main text or the supplementary materials.

## Supplementary Materials

Figs. S1 to S18

Tables S1 to S7

