## Supplementary Information for "Mapping the allosteric effects that define functional activity of SARS-CoV-2 specific antibodies"

**This file includes:**

Figs. S1 to S18

Tables S1 to S4

### Supplementary Figures

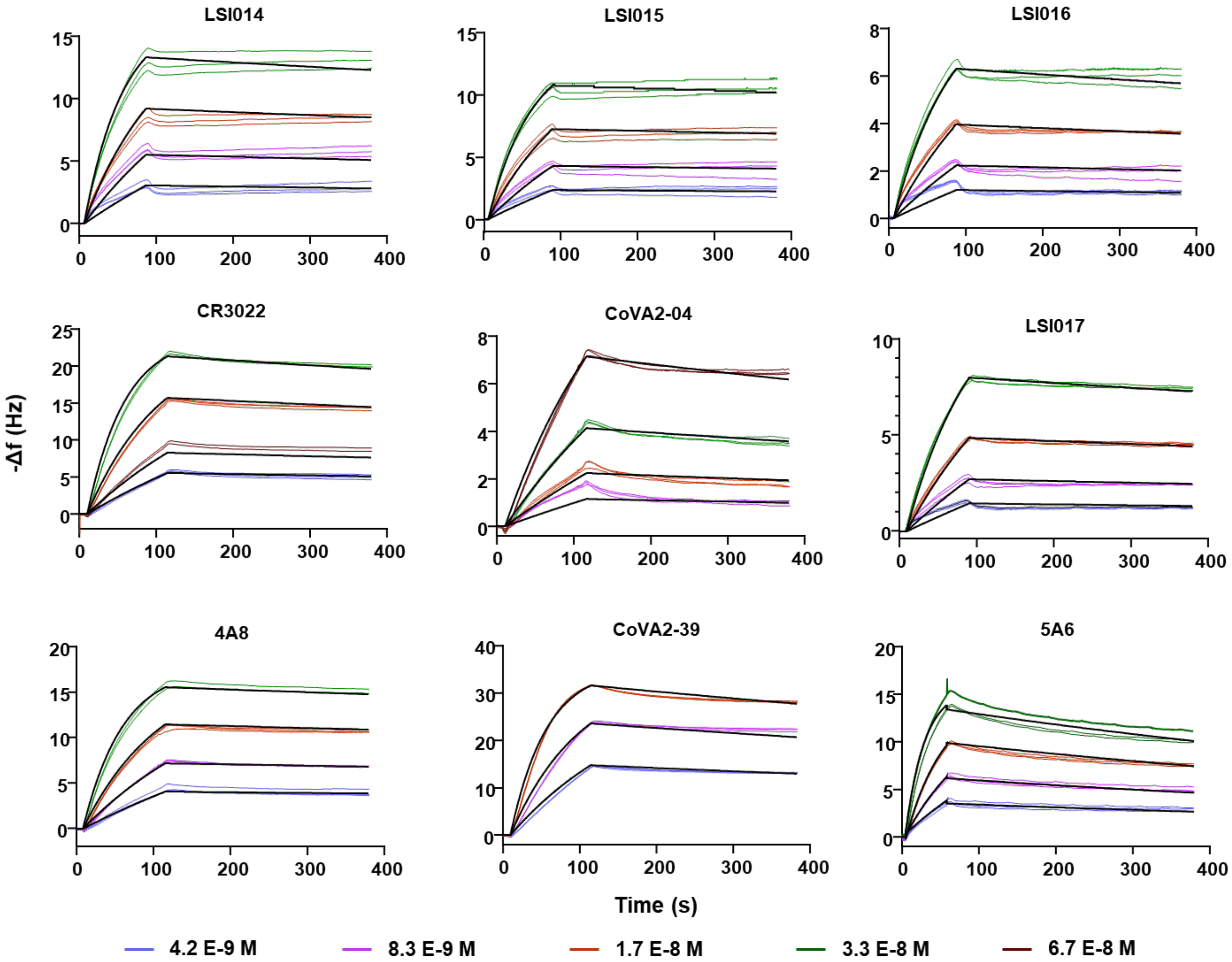


**Figure S1. QCM kinetic evaluation of the interactions between Spike and SARS CoV-2 antibodies.** Kinetic plots showing association-dissociation of in-house (LSI-CoVA-014, LSI-CoVA-015, LSI-CoVA-016, and LSI-CoVA-017) and previously studied (CR3022, CoVA2-04, 4A8, CoVA2-39, and 5A6) huMAbs. Antibodies at varying concentrations (4.2 nM – 66.7 nM) were flowed over Spike trimer immobilized onto the surface of a vibrating quartz crystal. Each experiment was performed in triplicate. Thick black lines show the theoretical 1:1 fit obtained using TraceDrawer evaluation software (Ridgeview Instruments).

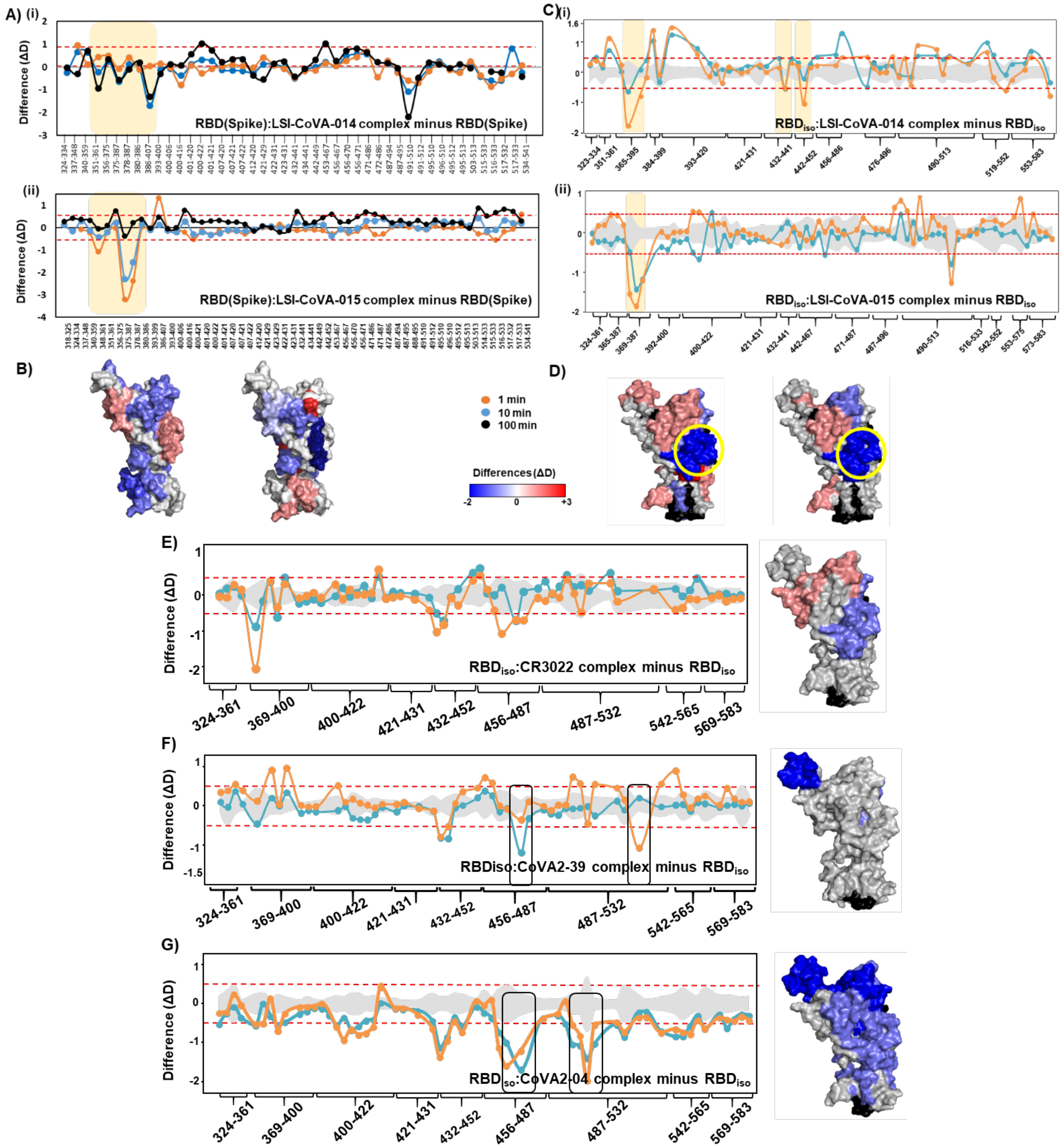


**Figure S2: HDXMS reveals antibodies recognizing different epitopes on RBD.**

Plots showing differences in deuterium exchange for HDX analysis of RBD of Spike with (A) LSI-CoVA-014 and (C) LSI-CoVA-015, and $RBD_{iso}$ with (D) CR3022, (E) CoVA2-04, and (F) CoVA2-39 with free RBD are shown for each peptide. Overlapping peptides were grouped and their residue numbers indicated (Tables S6 and S7). Deuterium exchange was carried out at indicated labelling times, with positive differences indicating increased exchange, and negative differences denoting decreased deuterium exchange in the antibody-bound complex. Cryptic epitope sites are highlighted in yellow (A-D), while RBM epitope sites are highlighted in black box (F-G). A significance threshold of ± 0.5 D is indicated by red-dashed lines with deviations in grey. Deuterium exchange differences at 10 min labelling are mapped on to structure (surface representation) of RBD, as per key.

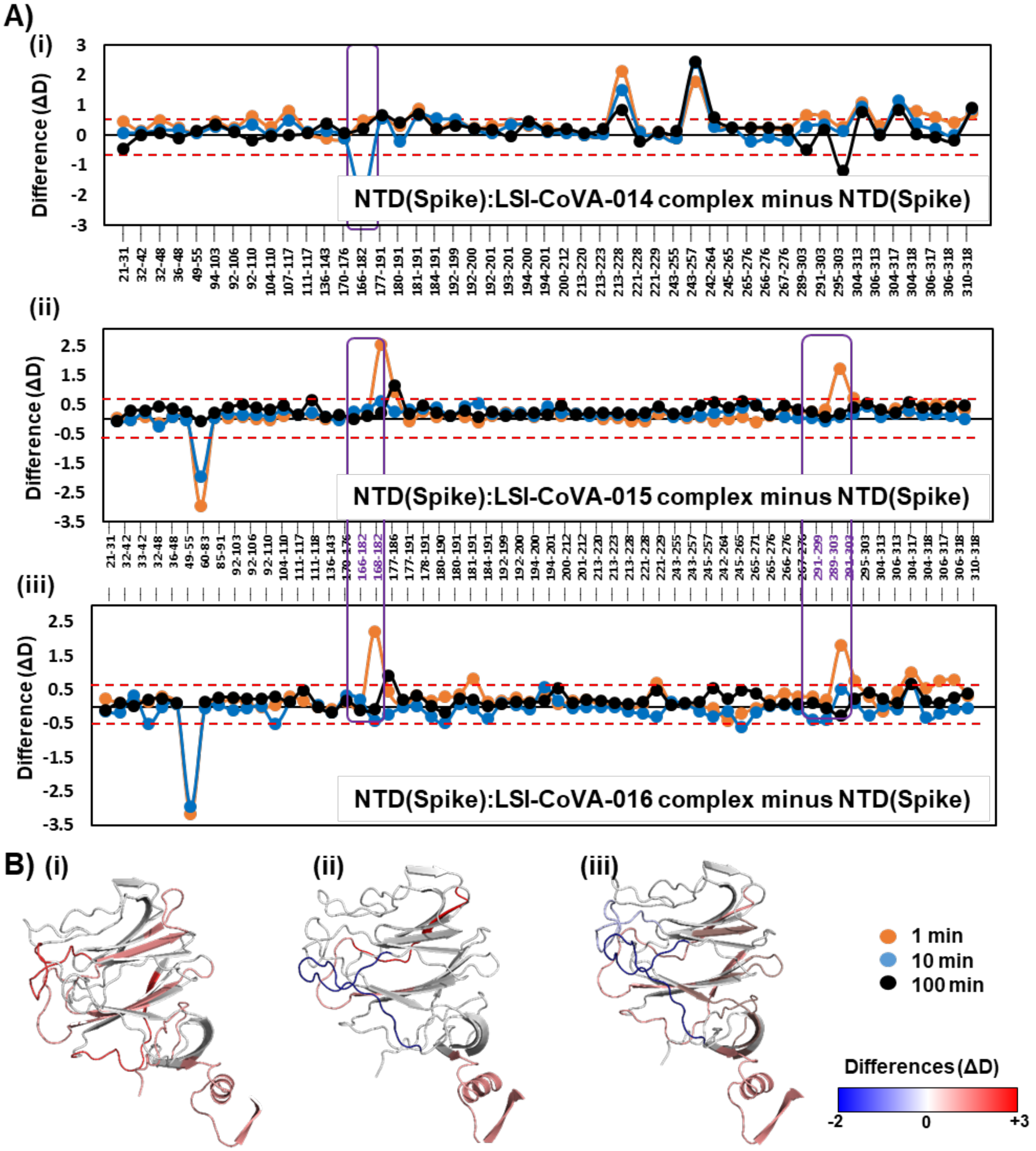


**Figure S3: Determining the effect of RBD-binding antibodies on NTD of the S1 subunit.**
Plots of differences in deuterium exchange between antibody-bound and free states of NTD of Spike trimer, with the residue numbers indicated as per Table S6. (A) Differences upon binding of (i) LSI-CoVA-014, (ii) LSI-CoVA-015, and (iii) LSI-CoVA-016 to the Spike trimer are shown. Purple boxes highlighted peptides of NTD interacting with RBD (166-182) and the S1 subunit (289-305). A threshold of ± 0.5 D considered significant is highlighted by red-dashed lines, and standard deviation is in grey. (B) Differences in deuterium exchange at 1 min labeling time were mapped onto structures of NTD (Spike), as indicated.

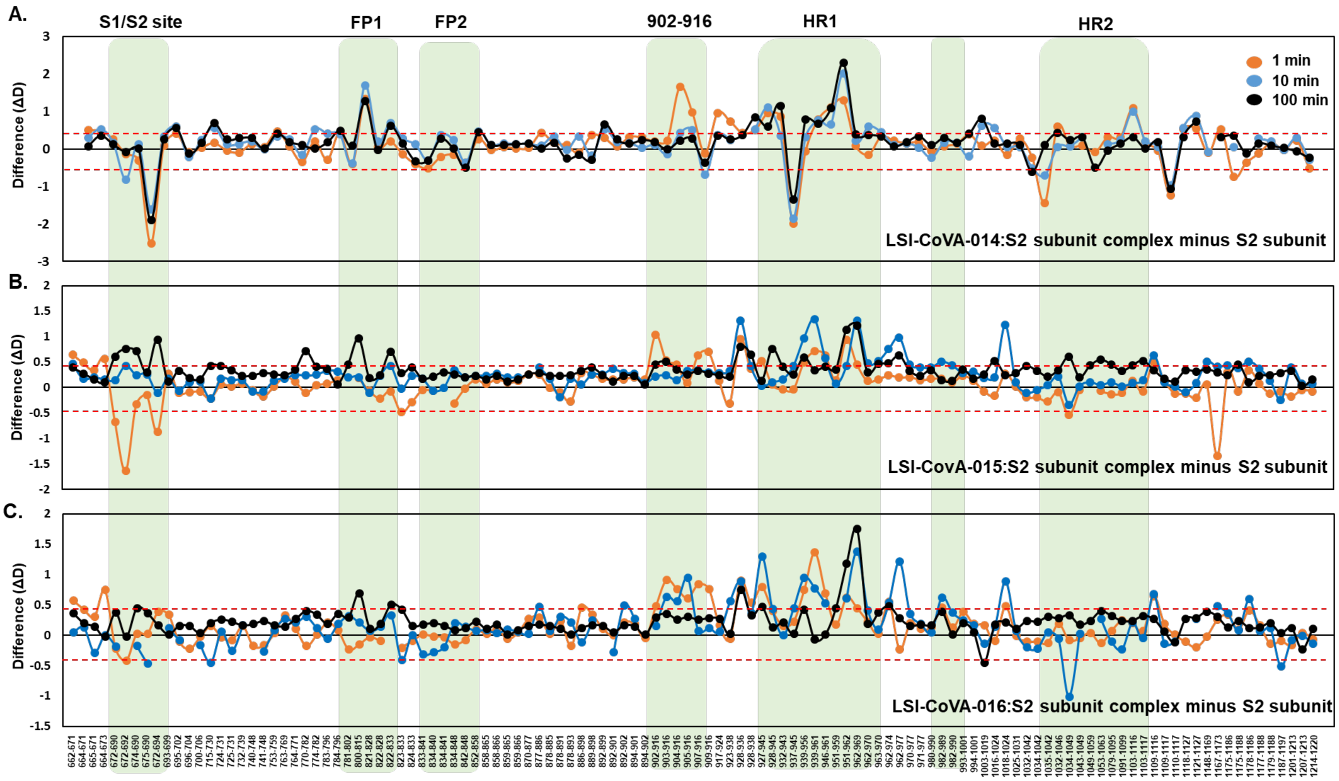


**Figure S4: Monitoring conformational changes induced at the S2 subunit by RBD-binding antibodies.**

Plots of differences in deuterium exchange between antibody-bound and free states of the S2 subunit of Spike trimer complexes with (A) LSI-CoVA-014, (B) LSI-CoVA-015, and (C) LSI-CoVA-016 are shown for indicated labeling times. Residue numbers are indicated for pepsin-proteolyzed peptides of the S2 subunit (Table S6), with important regions (S1/S2 cleavage site, Fusion peptide 1 and 2, Heptad Repeats 1 and 2, Central helix) highlighted in green.

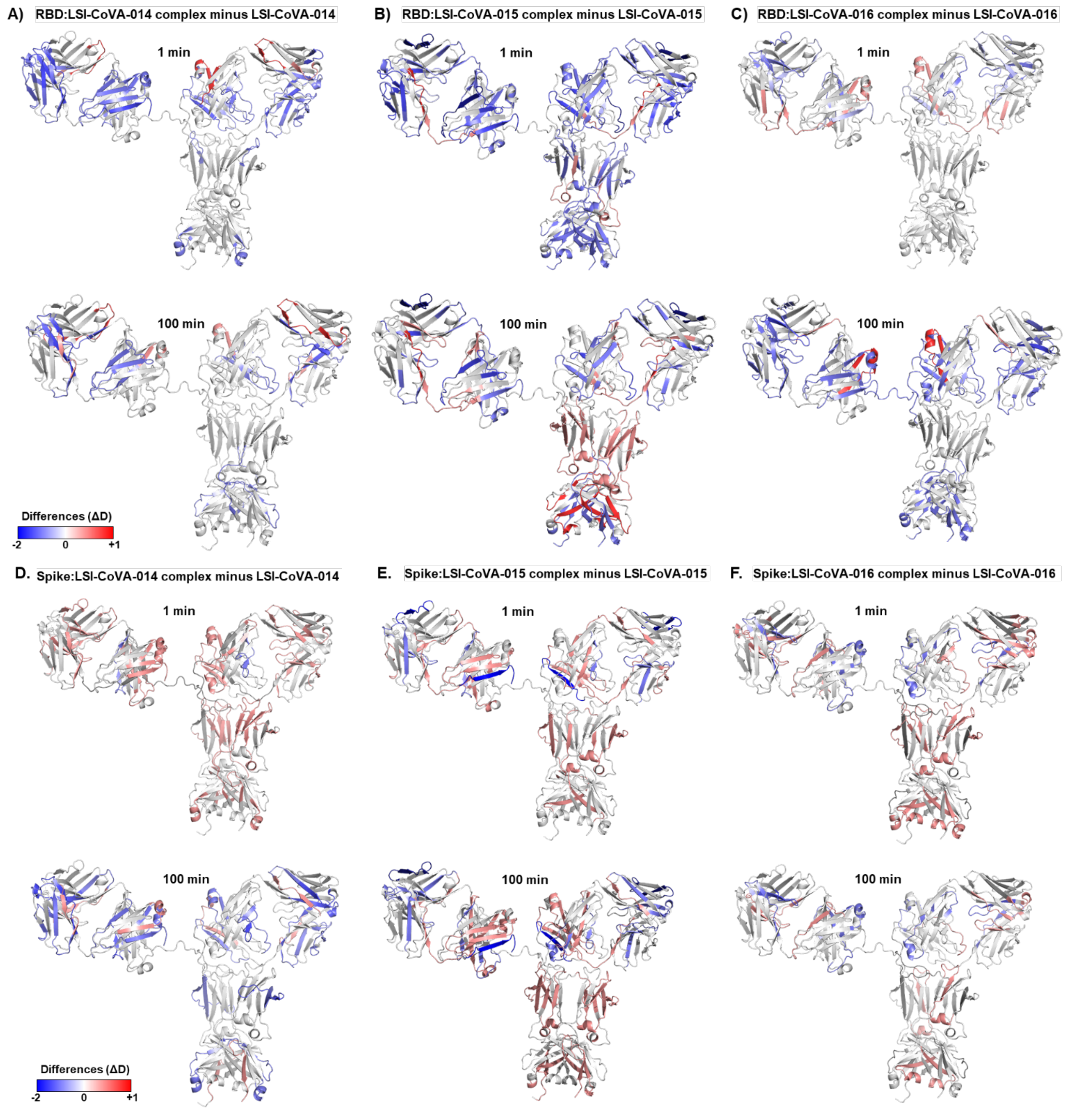


**Figure S5. Probing changes at the paratope sites of LSI-CoVA-014, LSI-CoVA-015 and LSI-CoVA-016.**

Cartoon representation of a model of antibody showing the differences in deuterium exchange between $RBD_{iso}$-bound and free states of (A) LSI-CoVA-014, (B) LSI-CoVA-015, and (C) LSI-CoVA-016 and Spike-bound and free states of (D) LSI-CoVA-014, (E) LSI-CoVA-015, and (F) LSI-CoVA-016 at 1 (top) and 100 (bottom) min labeling timepoints, as indicated. Changes in deuterium exchange at the paratope-sites of light (CDRL1-L3) and heavy (CDRH1-H3) chains of each antibody are tabulated in Table S2.

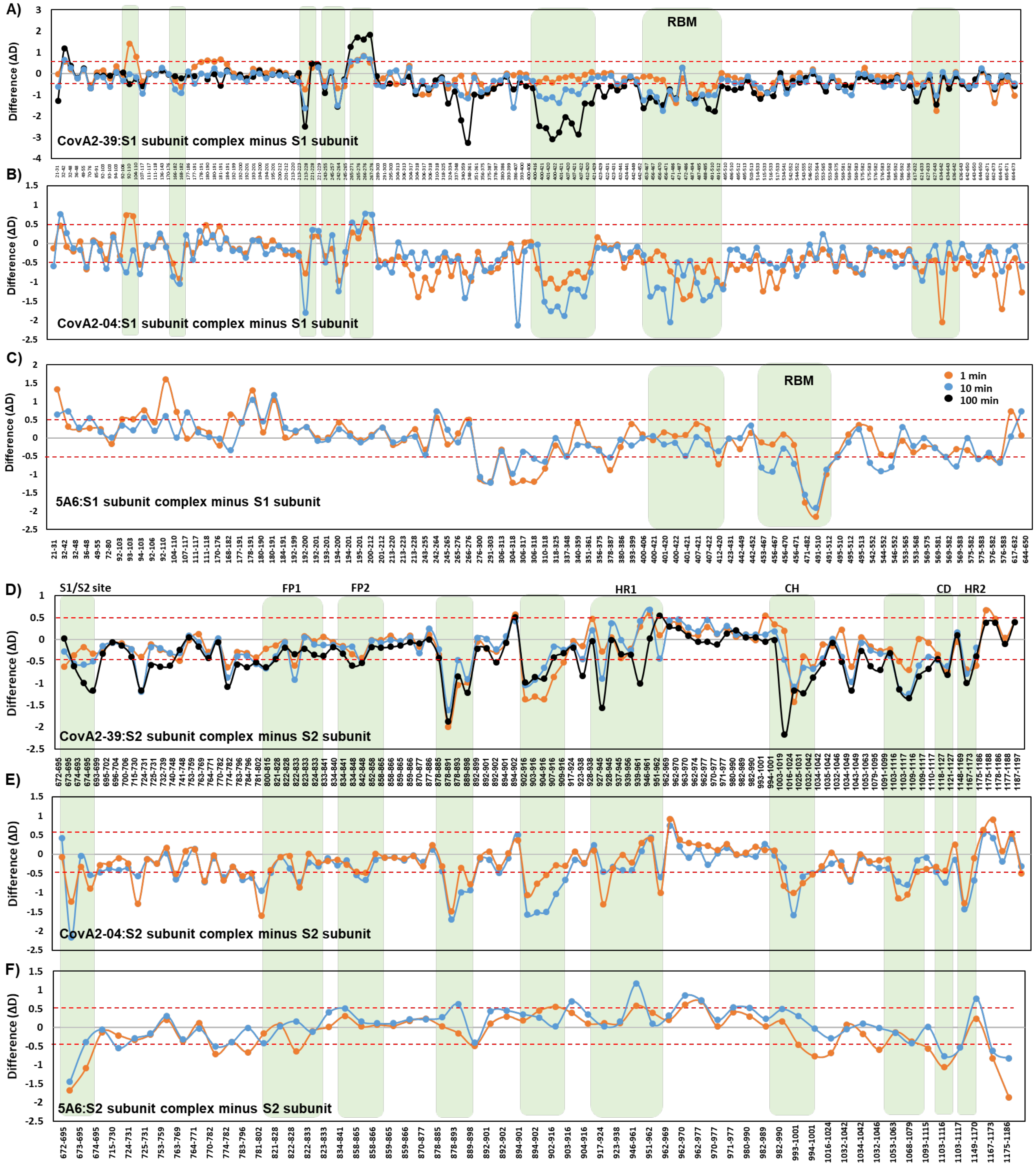


**Figure S6: RBM binding antibodies stabilize the Spike dynamics.**

Plots showing differences in deuterium exchange kinetics of the S1 (A-C) and the S2 (D-F) subunits of Spike in the presence and absence of (A, D) CoVA2-39, (B, E) CoVA2-04, and (C, F) 5A6 HuMAbs for various pepsin-digest fragments and their residue numbers indicated as per Table S6. A threshold of ± 0.5 D is considered significant and indicated by red-dashed line. Green boxes highlight the epitope sites spanning RBM, and important sites of the S2 subunit.

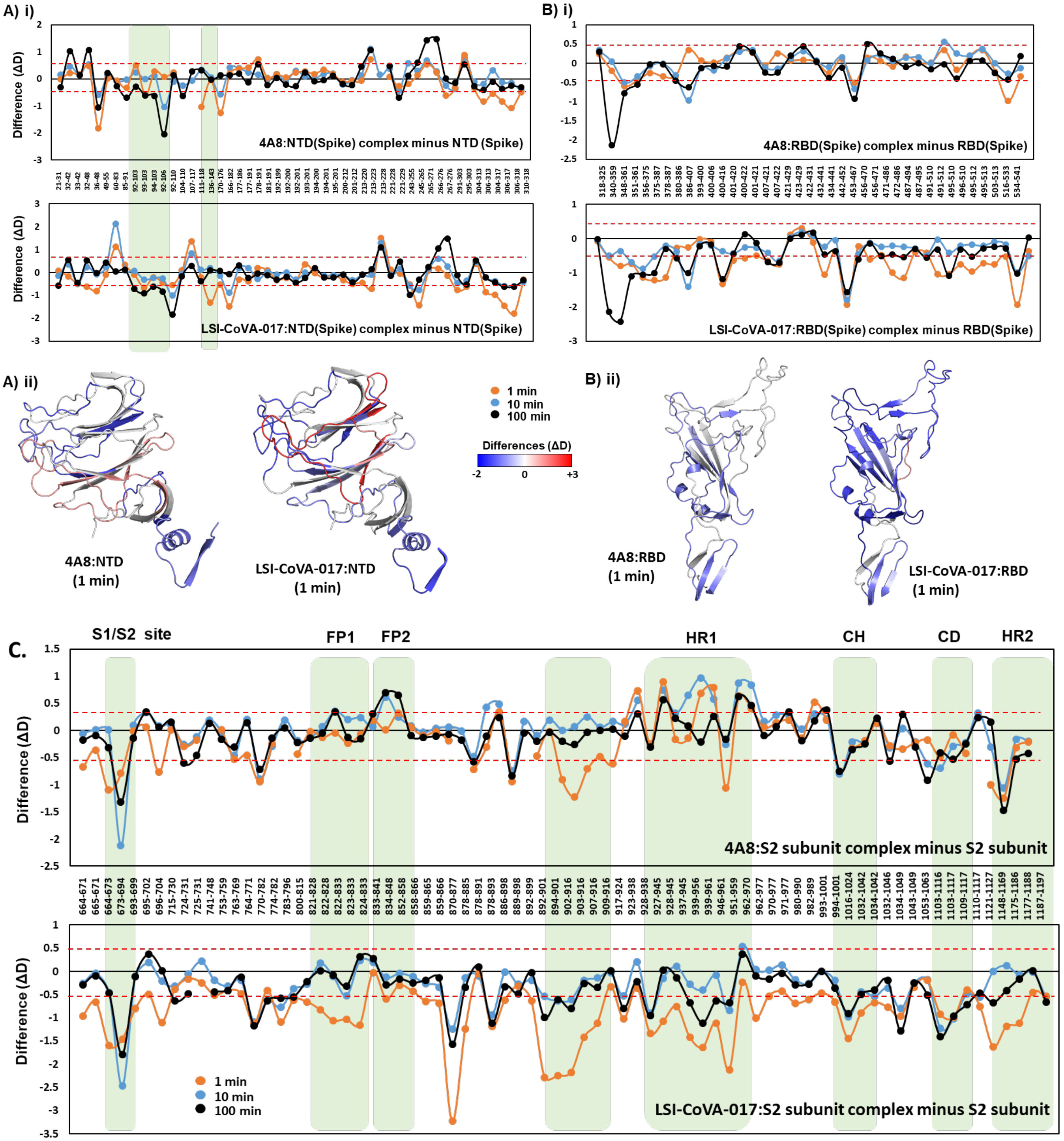


**Figure S7: Determining the effect of NTD-binding antibodies on S1 and S2 subunits.**
Plots of differences in deuterium exchange for Spike bound to 4A8- (top panels) and LSI-CoVA-017 (bottom panels) antibodies are shown. For clarity, peptides spanning (A) NTD, (B) RBD, and (C) S2 subunit are indicated with their residue numbers as per Table S6. A significance threshold of ± 0.5D was considered and indicated by red-dashed lines. Green boxes highlight important regions. Differences in deuterium exchange values at 1 min labeling timepoint were mapped onto structures of Spike, with close-up views of (A ii) NTD and (B ii) RBD shown for clarity.

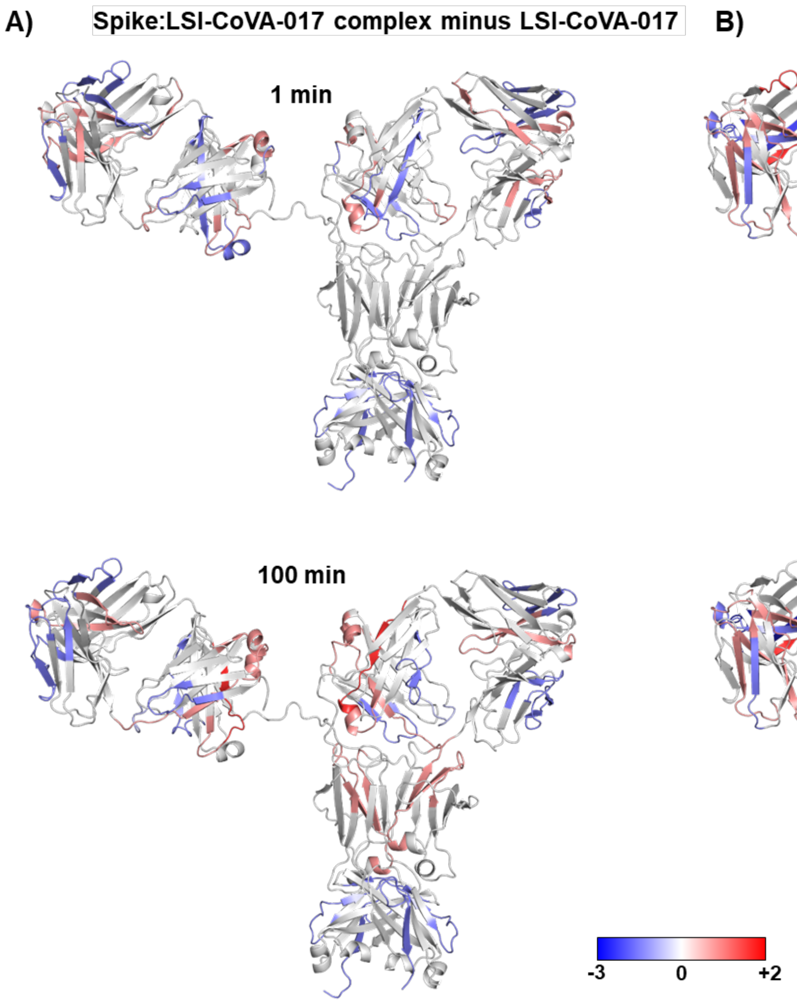


**Figure S8. Monitoring the effects of Spike on NTD-binding antibodies LSI-CoVA-017 and 4A8.** Cartoon representation of a model of antibody showing the differences in deuterium exchange between Spike-bound and free states of the NTD-binding antibody (A) LSI-CoVA-017 (novel antibody in this study), in comparison with (B) 4A8 antibody (previously characterized) at 1 and 100 min labeling timepoints, as indicated. Changes in deuterium exchange at the paratope-sites of light (CDRL1-L3) and heavy (CDRH1-H3) chains of each antibody are tabulated in Table 1.

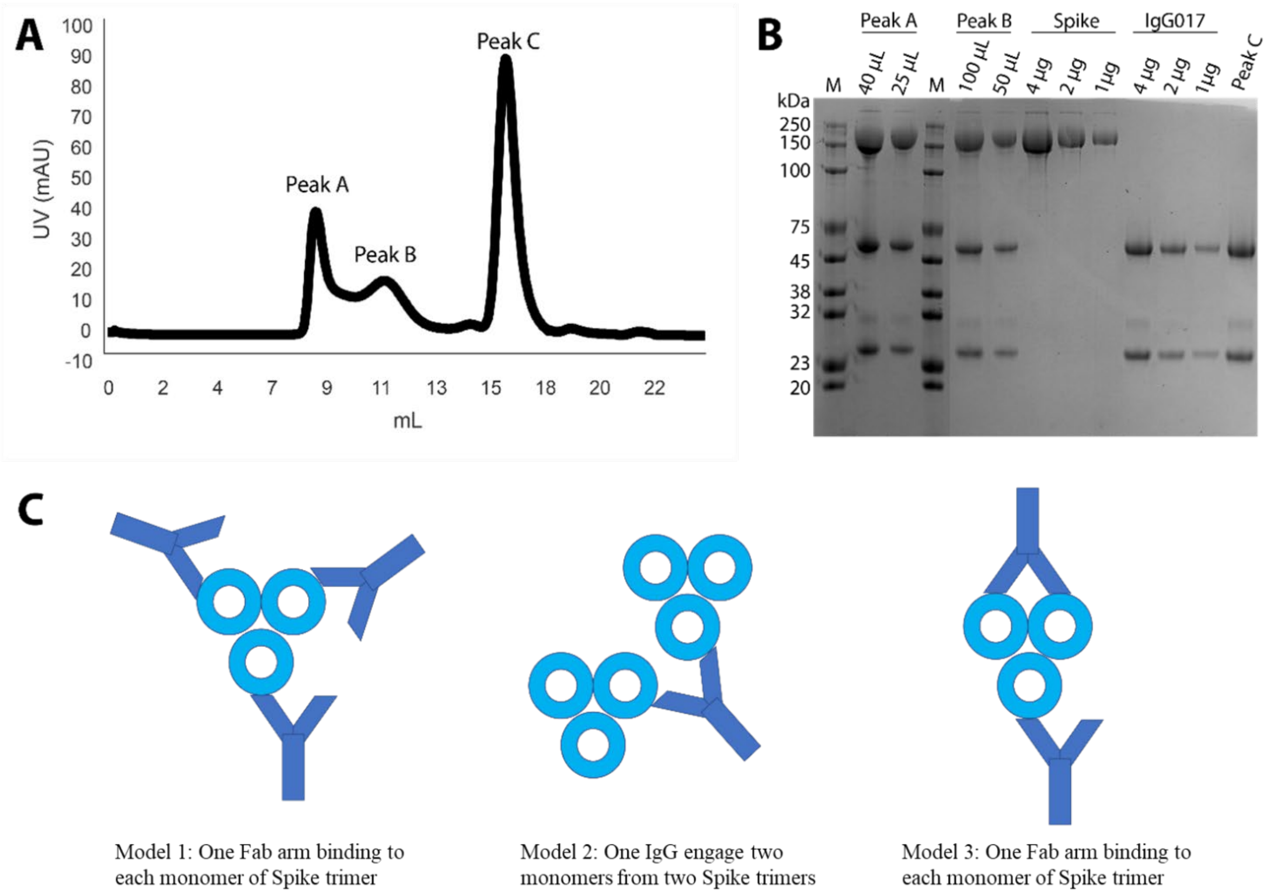


**Figure S9. Characterization of Spike:LSI-CoVA-017 interaction by size-exclusion chromatography and SDS-PAGE densitometry.**

(A) Chromatogram of Spike:LSI-CoVA-017 mixture following injection on a Superose 6 Increase 10/300 GL gel filtration column. Peak A corresponds to high molecular weight oligomers, peak B contains lower molecular weight oligomers of Spike:LSI-CoVA-017 complexes, and peak C corresponds to unbound LSI-CoVA-017 added in excess. (B) Image of denaturing polyacrylamide electrophoretic analysis of peak fractions of A and B. LSI-CoVA-017 and Spike were loaded for reference and for densitometry analysis calibration. (C) Schematics of the three most plausible binding modes of interaction between LSI-CoVA-017 and Spike trimer, based on the stoichiometric ratios determined and listed in Table S4.

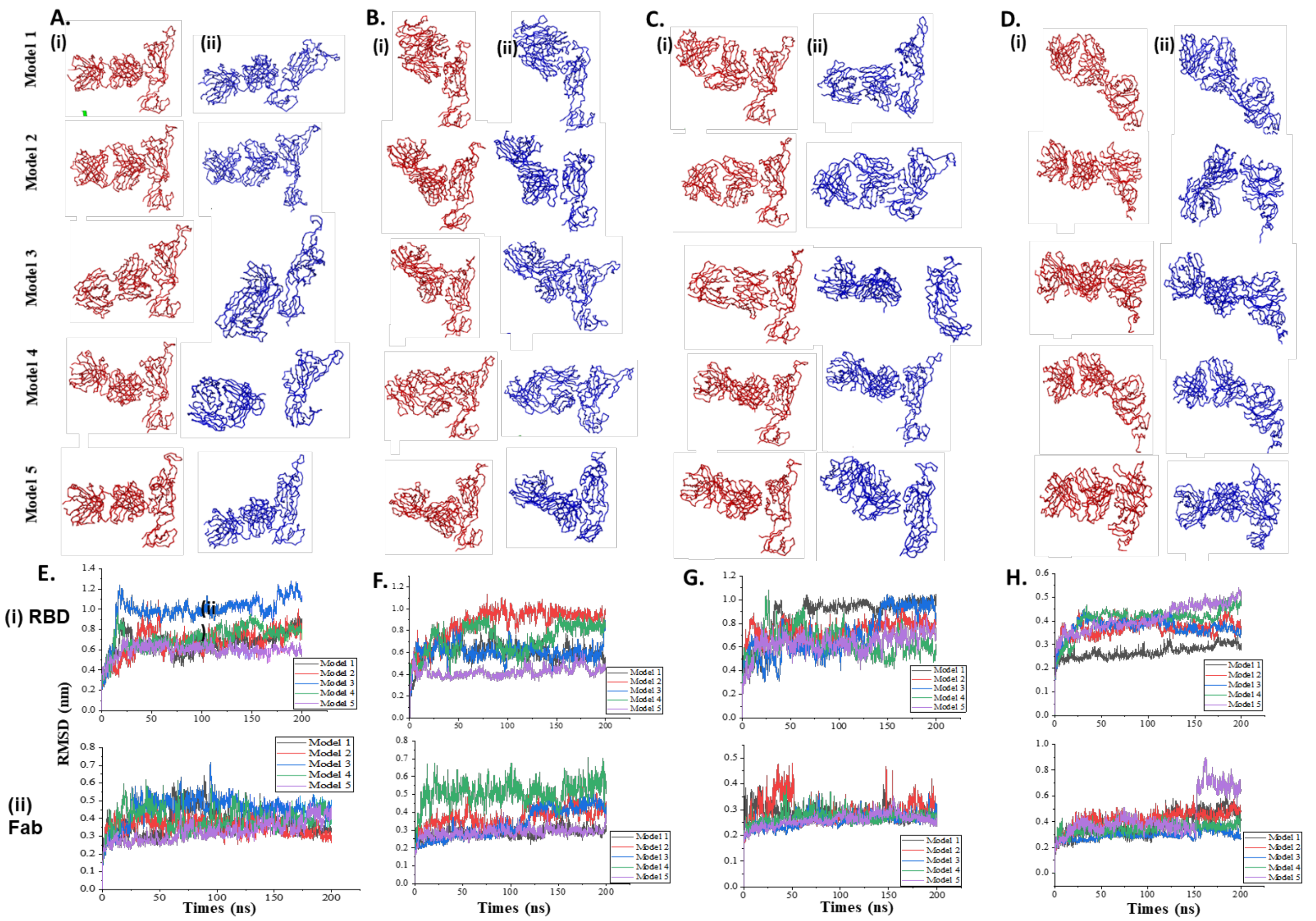


**Figure S10: Visualizing the orientation of** Fab:RBD/NTD **complexes.**

(A) LSI-CoVA-014:RBD, (B) LSI-CoVA-015:RBD, (C) LSI-CoVA-016:RBD, and (D) LSI-CoVA-017:NTD Fab-antigen complexes in trace representation at the beginning (red) and the end (blue) of a 200 ns simulation. (E-H) Backbone RMSD of Fab, RBD and NTD after least-squares fit to the backbone of the complexes for five models from the four systems for 200 ns long simulations.

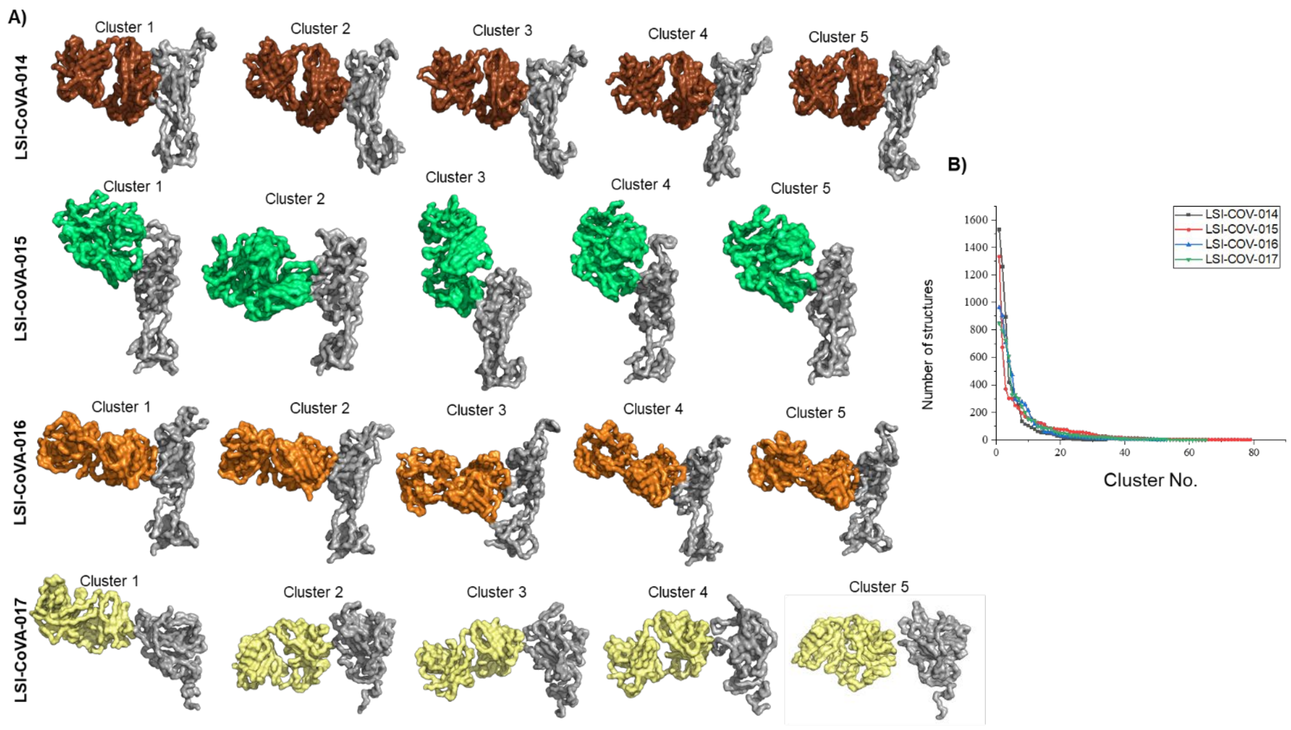


**Figure S11. Cluster analysis identifies the dominant orientations of Fab:RBD/NTD models.** A) Surface representation of central structures of 5 most populated clusters from LSI-CoVA-014:RBD, LSI-CoVA-015:RBD, LSI-CoVA-016:RBD, LSI-CoVA-017:NTD complexes. B) Plot showing the size of each cluster of RBD with LSI-CoVA-014 (black), LSI-CoVA-015 (red), LSI-CoVA-016 (cyan), and LSI-CoVA-017 (green) using GROMOS method with an RMSD cut-off of 0.35 nm.

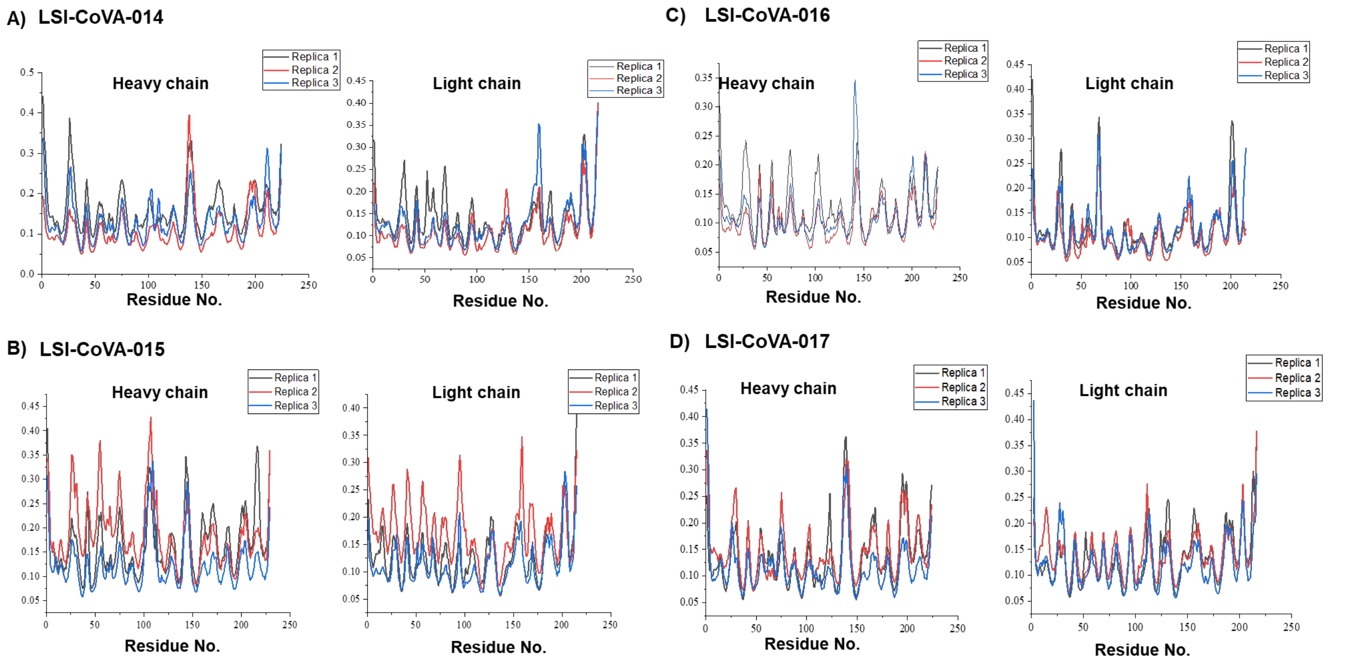


**Figure S12: Determining the variations of simulations between different Fab:RBD/NTD models.** Root mean square fluctuation (RMSF) of triplicate simulations of (A) LSI-CoVA-014, (B) LSI-CoVA-015, (C) LSI-CoVA-016, and (D) LSI-CoVA-017 antibodies for heavy (left) and light (right) chains from each model bound to RBD and NTD.

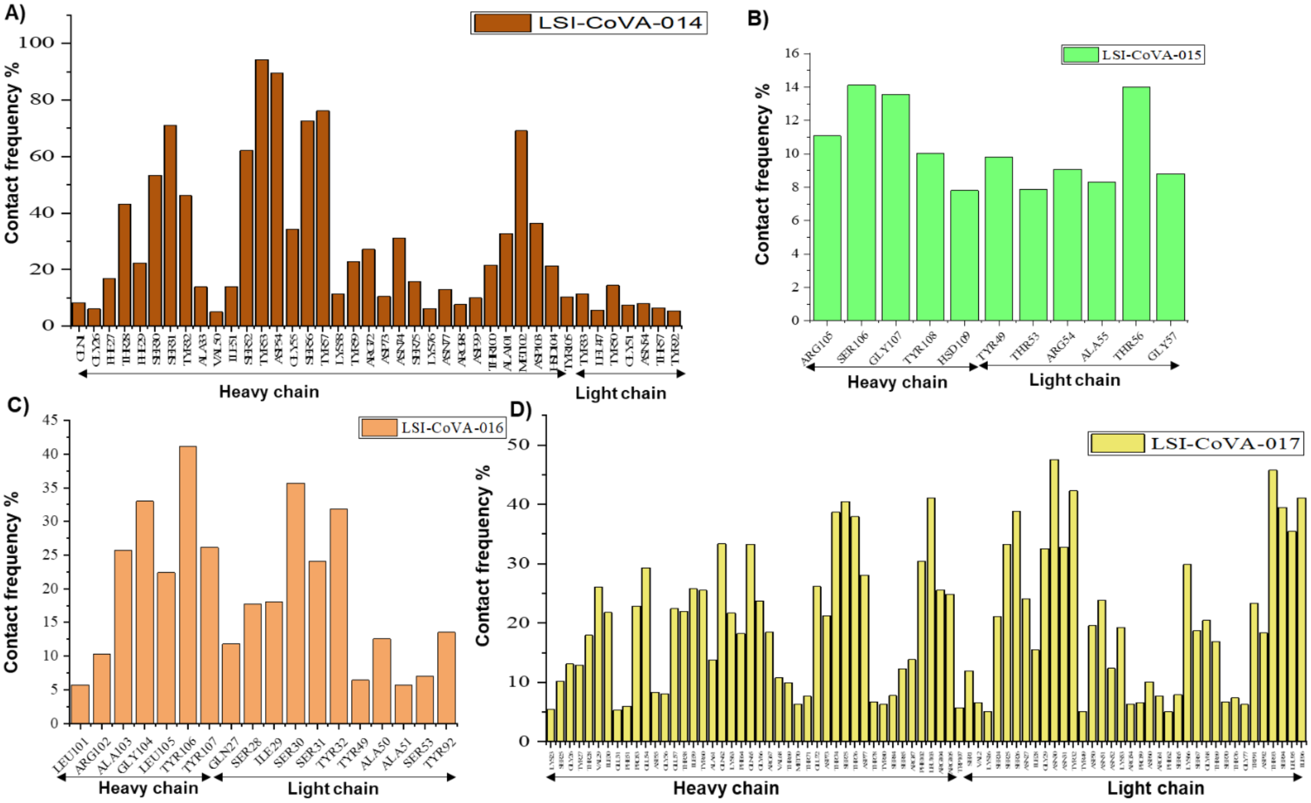


**Figure S13: Interactions of the glycan moiety with the Fab molecule.**

Contact frequencies % greater than 0.1 (residues interacting with glycan for at least 10% of simulation time) of the Fab residues from (A) LSI-CoVA-014, (B) LSI-CoVA-015, (C) LSI-CoVA-016, and (D) LSI-CoVA-017 with the glycan moiety of RBD or NTD during 200 ns of simulation time. Models wherein Fab separates from RBD or NTD were not considered and the maximum contact frequency value of each residue from remaining models is plotted.

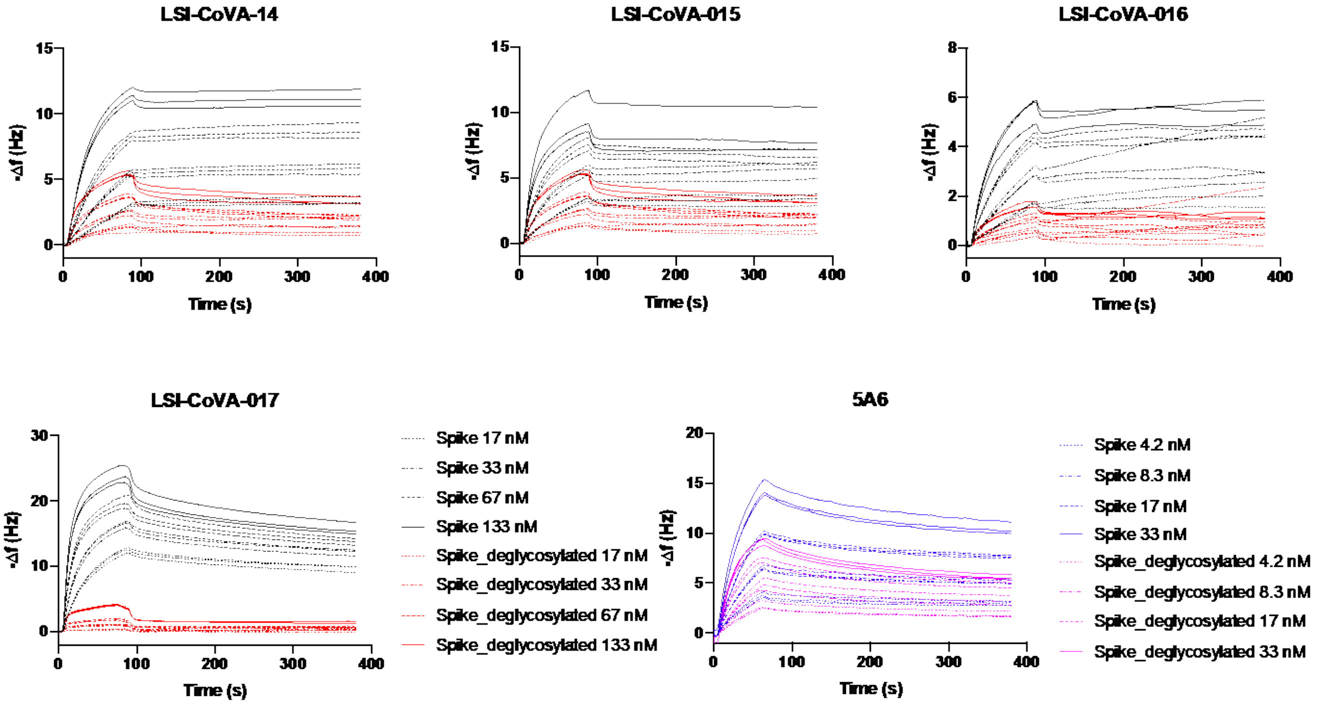


**Figure S14: Binding of novel antibodies and 5A6 to glycosylated and deglycosylated Spike trimer.** Antibodies at varying concentrations (4.2 nM – 133 nM) were flowed over Spike/deglycosylated Spike trimer immobilized onto the surface of a vibrating quartz crystal. Each experiment was performed in duplicate or triplicate.

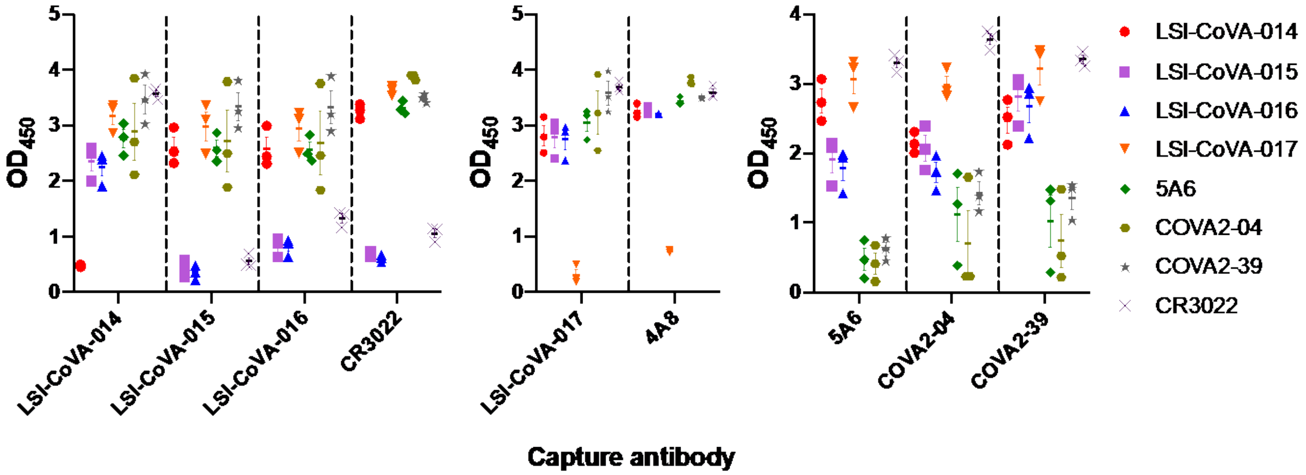


**Figure S15. Capture ELISA for pairs of selected antibodies**

SARS-CoV-2 Spike (0.1 μg) was captured by LSI-CoVA-014, LSI-CoVA-015, LSI-CoVA-016 and CR3022 (A); LSI-CoVA-017 and 4A8 (B) and 5A6, COVA2-04 and COVA2-39 (C), detected using peroxidase-labelled monoclonal antibodies. A low OD450 value is indicative of impaired binding of the peroxidase-labelled detection antibody, as listed. Data was collected from three individual experiments and represented as mean ± S.E.M.

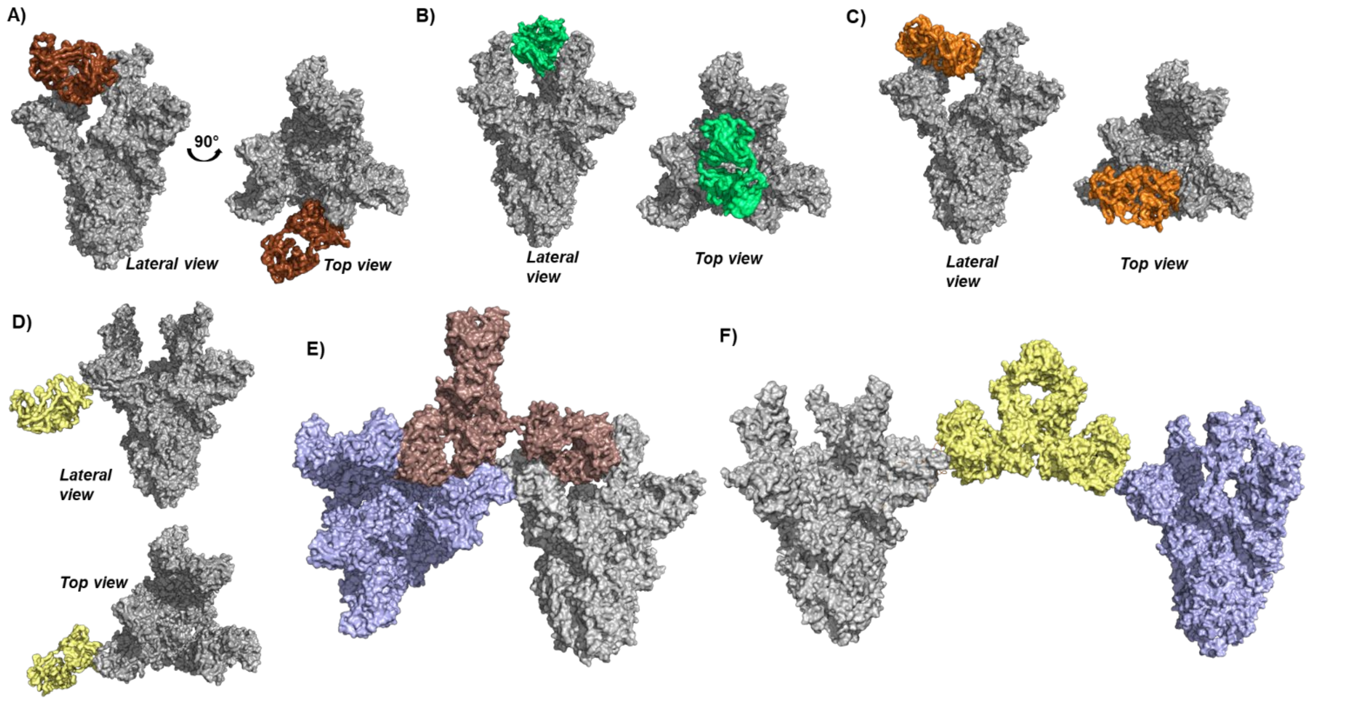


**Figure S16. Models of Spike:antibody complexes.**
Lateral and side views of surface representations showing Fab arms of (A) LSI-CoVA-014 (brown), (B) LSI-CoVA-015 (green), (C) LSI-CoVA-016 (orange), and (D) LSI-CoVA-017 (yellow) docked onto the Spike trimer (grey, PDB: 7A98). Representative structures from most populated cluster ('cluster 1') of Fab:RBD and Fab:NTD cluster analysis respectively was used. Surface representation of a model showing two Spike trimers (grey and blue) bound to both Fab arms of (E) LSI-CoVA-014 (brown) and (F) LSI-CoVA-017 (yellow) are predicted.

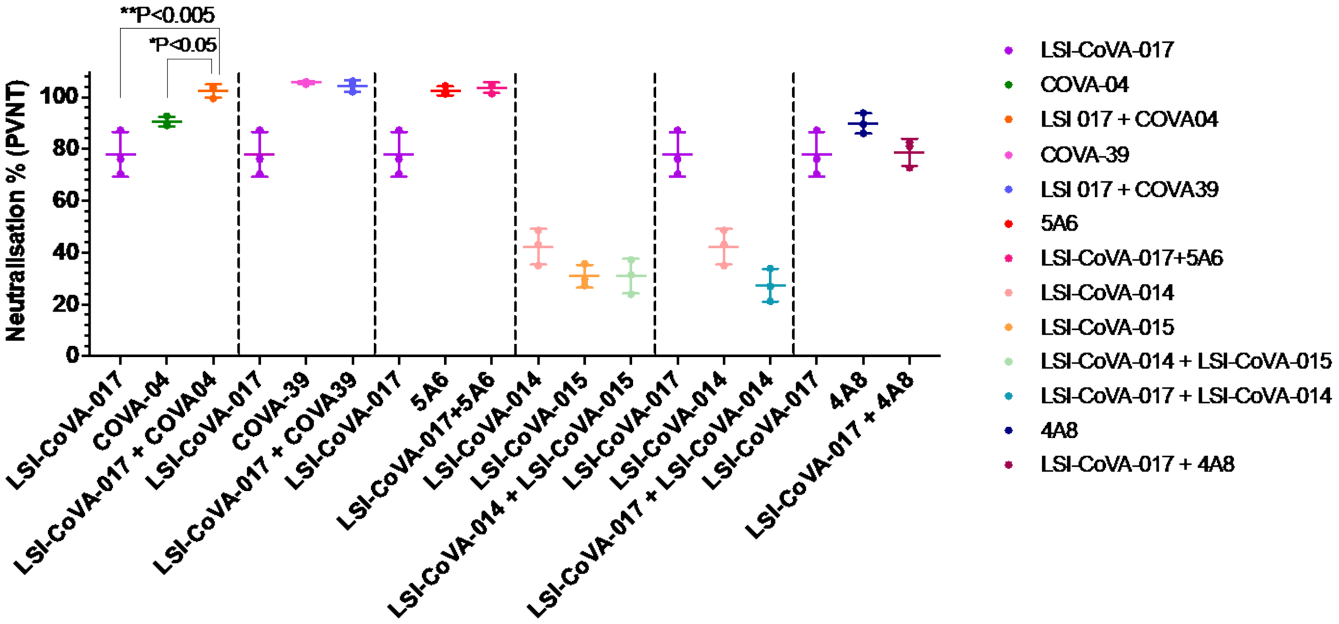


**Figure S17. Neutralization of pseudo SARS-CoV-2 virus using antibody cocktail.**
Pair-mAb cocktail in the ration of 1:9 to a final total concentration of 10 µg/ml or the single huMAb at a concentration of 10 µg/ml were incubated with pseudovirus lentiviral construct expressing the SARS-CoV-2 Spike protein. The antibody:pseudovirus mixtures were then added to CHO-ACE2 cells. The chemiluminescence readout from the luciferase-tagged reporter in the lentiviral construct, was then plotted and represented as percentage neutralization. The data is shown as mean ± s.d of triplicate measurements. The plots and one-way ANOVA statistical analysis were done using Graphpad prism.

M F V F L V L L P L V S S Q C V N L T T R T Q L P P A Y T N S F T R G V Y Y P D K V F R S S V L H S T Q D L F L P F F S
5 10 15 20 25 30 35 40 45 50 55 60

N V T W F H A I H V S G T N G T K R F D N P V L P F N D G V Y F A S T E K S N I I R G W I F G T T L D S K T Q S L L I V
65 70 75 80 85 90 95 100 105 110 115 120

N N A T N V V I K V C E F Q F C N D P F L G V Y Y H K N N K S W M E S E F R V Y S S A N N C T F E Y V S Q P F L M D L E
125 130 135 140 145 150 155 160 165 170 175 180

G K Q G N F K N L R E F V F K N I D G Y F K I Y S K H T P I N L V R D L P Q G F S A L E P L V D L P I G I N I T R F Q T
185 190 195 200 205 210 215 220 225 230 235 240

L L A L H R S Y L T P G D S S S G W T A G A A A Y Y V G Y L Q P R T F L L K Y N E N G T I T D A V D C A L D P L S E T K
245 250 255 260 265 270 275 280 285 290 295 300

C T L K S F T V E K G I Y Q T S N F R V Q P T E S I V R F P N I T N L C P F G E V F N A T R F A S V Y A W N R K R I S N
305 310 315 320 325 330 335 340 345 350 355 360

C V A D Y S V L Y N S A S F S T F K C Y G V S P T K L N D L C F T N V Y A D S F V I R G D E V R Q I A P G Q T G K I A D
365 370 375 380 385 390 395 400 405 410 415 420

Y N Y K L P D D F T G C V I A W N S N N L D S K V G G N Y N Y L Y R L F R K S N L K P F E R D I S T E I Y Q A G S T P C
425 430 435 440 445 450 455 460 465 470 475 480

N G V E G F N C Y F P L Q S Y G F Q P T N G V G Y Q P Y R V V V L S F E L L H A P A T V C G P K K S T N L V K N K C V N
485 490 495 500 505 510 515 520 525 530 535 540

F N F N G L T G T G V L T E S N K K F L P F Q Q F G R D I A D T T D A V R D P Q T L E I L D I T P C S F G G V S V I T P
545 550 555 560 565 570 575 580 585 590 595 600

G T N T S N Q V A V L Y Q D V N C T E V P V A I H A D Q L T P T W R V Y S T G S N V F Q T R A G C L I G A E H V N N S Y
605 610 615 620 625 630 635 640 645 650 655 660

E C D I P I G A G I C A S Y Q T Q T N S P G S A S S V A S Q S I I A Y T M S L G A E N S V A Y S N N S I A I P T N F T I
665 670 675 680 685 690 695 700 705 710 715 720

S V T T E I L P V S M T K T S V D C T M Y I C G D S T E C S N L L L Q Y G S F C T Q L N R A L T G I A V E Q D K N T Q E
725 730 735 740 745 750 755 760 765 770 775 780

V F A Q V K Q I Y K T P P I K D F G G F N F S Q I L P D P S K P S K R S F I E D L L F N K V T L A D A G F I K Q Y G D C
785 790 795 800 805 810 815 820 825 830 835 840

L G D I A A R D L I C A Q K F N G L T V L P P L L T D E M I A Q Y T S A L L A G T I T S G W T F G A G A A L Q I P F A M
845 850 855 860 865 870 875 880 885 890 895 900

Q M A Y R F N G I G V T Q N V L Y E N Q K L I A N Q F N S A I G K I Q D S L S S T A S A L G K L Q D V V N Q N A Q A L N
905 910 915 920 925 930 935 940 945 950 955 960

T L V K Q L S S N F G A I S S V L N D I L S R L D P P E A E V Q I D R L I T G R L Q S L Q T Y V T Q Q L I R A A E I R A
965 970 975 980 985 990 995 1000 1005 1010 1015 1020

S A N L A A T K M S E C V L G Q S K R V D F C G K G Y H L M S F P Q S A P H G V V F L H V T Y V P A Q E K N F T T A P A
1025 1030 1035 1040 1045 1050 1055 1060 1065 1070 1075 1080

I C H D G K A H F P R E G V F V S N G T H W F V T Q R N F Y E P Q I I T T D N T F V S G N C D V V I G I V N N T V Y D P
1085 1090 1095 1100 1105 1110 1115 1120 1125 1130 1135 1140

L Q P E L D S F K E E L D K Y F K N H T S P D V D L G D I S G I N A S V V N I Q K E I D R L N E V A K N L N E S L I D L
1145 1150 1155 1160 1165 1170 1175 1180 1185 1190 1195 1200

Q E L G K Y E Q V D G
1205 1210

Total: 267 Peptides, 85.4% Coverage, 2.98 Redundancy

**Figure S18: Pepsin-digest map of Spike.**
HDXMS analysis resulted in total of 267 pepsin-digested fragments (‘peptides’) spanning 86% of the Spike protein. Each peptide is indicated by green line. The sequence shown is for the Spike (1-1208) construct used in this study.

| **Antibody** | $\mathbf{B_{max}}$ | $\mathbf{k_a}$ **(1/[(M-s)]** | $\mathbf{k_d}$ **[1/s]** | $\mathbf{K_D}$ **(M)** | **Chi²** |
|---|---|---|---|---|---|
| LSI-CoVA-016 | 9.61 | 3.90 E+05 | 3.00 E-04 | 7.68 E-10 | 0.06 |
| LSI-CoVA-017 | 19.66 | 4.46 E+05 | 7.48 E-05 | 1.67 E-10 | 0.12 |
| 4A8 | 18.03 | 5.74 E+05 | 1.95 E-04 | 3.40 E-10 | 0.10 |
| LSI-CoVA-015 | 13.86 | 5.61 E+05 | 1.88 E-04 | 3.35 E-10 | 0.21 |
| COVA2-39 | 36.07 | 1.25 E+06 | 4.87 E-04 | 3.88 E-10 | 0.57 |
| LSI-CoVA-014 | 17.12 | 5.57 E+05 | 2.88 E-04 | 5.16 E-10 | 0.35 |
| CR3022 | 24.53 | 6.05 E+05 | 3.16 E-04 | 5.21 E-10 | 0.38 |
| COVA2-04 | 15.25 | 9.32 E+04 | 5.55 E-04 | 5.96 E-09 | 0.05 |
| 5A6 | 15.52 | 1.18 E+06 | 8.87 E-04 | 7.55 E-10 | 0.15 |

**Table S1:** Quartz crystal microbalance analysis of antibody binding kinetics to Hexapro Spike trimer (0.03 μM). $k_a$ = association constant; $k_d$: dissociation constant

| | CDRL1 | CDRL2 | CDRL3 | CDRH1 | CDRH2 | CDRH3 |
|---|---|---|---|---|---|---|
| | **RBD** | **RBD** | **RBD** | **RBD** | **RBD** | **RBD** |
| **LSI-CoVA-014** | **-1.00** | **-0.32** | **-1.86** | **0.65** | **-1.74** | **-1.70** |
| **LSI-CoVA-015** | **-0.51** | **-1.73** | **-0.70** | **-1.08** | **-1.18** | **0.41** |
| **LSI-CoVA-016** | **-0.23** | **-0.41** | **-0.76** | **-0.25** | **-0.55** | **0.24** |
| **LSI-CoVA-017** | **0.35** | **-0.36** | **-0.05** | **-0.21** | **-0.45** | **-0.17** |

| | CDRL1 | CDRL2 | CDRL3 | CDRH1 | CDRH2 | CDRH3 |
|---|---|---|---|---|---|---|
| | **Spike** | **Spike** | **Spike** | **Spike** | **Spike** | **Spike** |
| **LSI-CoVA-014** | **-0.55** | **-0.52** | **-0.81** | **0.45** | **0.40** | **-1.01** |
| **LSI-CoVA-015** | **0.17** | **-1.88** | **-0.41** | **-0.89** | **-0.27** | **0.60** |
| **LSI-CoVA-016** | **-0.16** | **-0.27** | **-0.71** | **-0.25** | **-0.20** | **0.80** |
| **LSI-CoVA-017** | **0.32** | **0.80** | **0.31** | **0.43** | **-0.51** | **-0.26** |

**Table S2:** Differences in deuterium exchange values for complementarity-determining regions (CDR) of light (CDR L1-L3) and heavy (CDR H1-H3) chains of various antibodies in the presence and absence of $RBD_{iso}$ (top) and Spike (bottom) as indicated. Positive differences (>0.5 D) are highlighted in red, and negative differences (<-0.5 D) are in green.

| Peak A (40 µl) | mg | MW(kDa) | pmol | Spike trimer or IgG (pmol) | LSI-CoVA-017:Spike trimer stoichiometry |
|---|---|---|---|---|---|
| Spike | 3.02 | 160.00 | 18.88 | 6.29 | |
| Heavy Chain | 1.87 | 50.00 | 37.33 | 18.67 | 2.96 |
| Light Chain | 0.95 | 25.00 | 37.87 | 18.93 | 3.01 |
| **Peak A (25 µl)** | **mg** | **MW(kDa)** | **pmol** | **Spike trimer or IgG (pmol)** | **LSI-CoVA-017:Spike trimer stoichiometry** |
| Spike | 2.02 | 160.00 | 12.63 | 4.21 | |
| Heavy Chain | 1.23 | 50.00 | 24.53 | 12.27 | 2.91 |
| Light Chain | 0.61 | 25.00 | 24.53 | 12.27 | 2.91 |
| **Peak B (100 µl)** | **mg** | **MW(kDa)** | **pmol** | **Spike trimer or IgG (pmol)** | **LSI-CoVA-017:Spike trimer stoichiometry** |
| Spike | 2.53 | 160.00 | 15.81 | 5.27 | |
| Heavy Chain | 1.57 | 50.00 | 31.33 | 15.67 | 2.97 |
| Light Chain | 0.82 | 25.00 | 32.67 | 16.33 | 3.09 |
| **Peak B (50 µl)** | **mg** | **MW(kDa)** | **pmol** | **Spike trimer or IgG (pmol)** | **LSI-CoVA-017:Spike trimer stoichiometry** |
| Spike | 1.30 | 160.00 | 8.13 | 2.71 | |
| Heavy Chain | 0.81 | 50.00 | 16.13 | 8.07 | 2.98 |
| Light Chain | 0.41 | 25.00 | 16.53 | 8.27 | 3.05 |

**Table S3: Determination of Spike:LSI-CoVA-017 binding stoichiometry**.
The absolute quantities of Spike, heavy and light chains of LSI-CoVA-017 were estimated based on the SDS-PAGE band intensities relative to quantification standards. The quantities of LSI-CoVA-017 were calculated based on the quantities of either heavy chain or light chain. Both approaches resulted in similar quantities determined of LSI-CoVA-017, and thus consistently suggest a binding stoichiometry of three LSI-CoVA-017 bound per Spike trimer. To confirm the reliability of the densitometry, samples were loaded at two different amounts and analyzed from each peak of size-exclusion chromatogram.

| System | Model# | No. of water ions | No. of $Na^{+}$ ions | No. of $Cl^{-}$ ions | Box size nm × nm × nm | Production run (ns) |
|---|---|---|---|---|---|---|
| **LSI-CoVA-014-RBD** | Model 1 | 71807 | 203 | 212 | 13.4×13.4×13.4 | 200 |
| | **Model 2** | **64934** | **185** | **194** | **13×13×13** | **200 × 3** |
| | Model 3 | 87998 | 250 | 259 | 14.3×14.3×14.3 | 200 |
| | Model 4 | 73478 | 208 | 217 | 13.5×13.5×13.5 | 200 |
| | Model 5 | 77021 | 218 | 227 | 13.7×13.7×13.7 | 200 |
| **LSI-CoVA-015-RBD** | **Model 1** | **95807** | **273** | **283** | **14.7×14.7×14.7** | **200 × 3** |
| | Model 2 | 78446 | 223 | 233 | 13.8×13.8×13.8 | 200 |
| | Model 3 | 85899 | 245 | 255 | 14.2×14.2×14.2 | 200 |
| | Model 4 | 57277 | 162 | 172 | 12.5×12.5×12.5 | 200 |
| | Model 5 | 57277 | 163 | 173 | 12.5×12.5×12.5 | 200 |
| **LSI-CoVA-016-RBD** | Model 1 | 76917 | 218 | 227 | 13.7×13.7×13.7 | 200 |
| | Model 2 | 80876 | 229 | 238 | 13.9×13.9×13.9 | 200 |
| | Model 3 | 82268 | 234 | 243 | 14×14×14 | 200 |
| | **Model 4** | **85898** | **245** | **254** | **14.2×14.2×14.2** | **200 × 3** |
| | Model 5 | 85946 | 245 | 254 | 14.2×14.2×14.2 | 200 |
| **LSI-CoVA-017-NTD** | Model 1 | 129534 | 368 | 372 | 16.2×16.2×16.2 | 200 |
| | Model 2 | 110527 | 314 | 318 | 15.4×15.4×15.4 | 200 |
| | **Model 3** | **85427** | **243** | **247** | **14.2×14.2×14.2** | **200 × 3** |
| | Model 4 | 115354 | 327 | 331 | 15.6×15.6×15.6 | 200 |
| | Model 5 | 78027 | 222 | 226 | 13.8×13.8×13.8 | 200 |

**Table S4: Monitoring dynamics of antigen-antibody complexes.**
Details of simulation setup for molecular dynamics of top 5 models of complexes between RBD/NTD and Fab domains of LSI-CoVA-014, LSI-CoVA-015, LSI-CoVA-016 and LSI-CoVA-017, as described in materials and methods. Models highlighted in bold show simulations performed in triplicates.
